# Mapping Lung Cancer Epithelial-Mesenchymal Transition States and Trajectories with Single-Cell Resolution

**DOI:** 10.1101/570341

**Authors:** Loukia G. Karacosta, Benedict Anchang, Nikolaos Ignatiadis, Samuel C. Kimmey, Jalen A. Benson, Joseph B. Shrager, Robert Tibshirani, Sean C. Bendall, Sylvia K. Plevritis

## Abstract

Elucidating a continuum of epithelial-mesenchymal transition (EMT) and mesenchymal-epithelial transition (MET) states in clinical samples promises new insights in cancer progression and drug response. Using mass cytometry time-course analysis, we resolve lung cancer EMT states through TGFβ-treatment and identify through TGFβ-withdrawal, an MET state previously unrealized. We demonstrate significant differences between EMT and MET trajectories using a novel computational tool (TRACER) for reconstructing trajectories between cell states. Additionally, we construct a lung cancer reference map of EMT and MET states referred to as the EMT-MET STAte MaP (STAMP). Using a neural net algorithm, we project clinical samples onto the EMT-MET STAMP to characterize their phenotypic profile with single-cell resolution in terms of our *in vitro* EMT-MET analysis. In summary, we provide a framework that can be extended to phenotypically characterize clinical samples in the context of *in vitro* studies showing differential EMT-MET traits related to metastasis and drug sensitivity.

## INTRODUCTION

Malignant cells often hijack biological processes that are utilized by their normal cell counterparts^1^. One example is the epithelial-mesenchymal transition (EMT), a developmental program critical to embryogenesis and wound healing^2,3^. Upon EMT initiation, epithelial cells undergo dramatic biochemical and morphological changes, including loss of polarity, loosening of tight and adherent junctions and acquisition of migratory and stem-like traits^4^. EMT is a dynamic process and under specific conditions is reversible (mesenchymal-epithelial transition, MET), highlighting a phenotypic plasticity that has been observed in both normal and malignant cells^3^.

The clinical significance of EMT in cancer is well documented, although controversial^5^. Mesenchymal signatures are indicators of poor prognosis in various adenocarcinomas, including lung^6,7^. In addition, several studies have demonstrated the role of EMT in drug response and resistance^8,9^ by offering evidence that EMT signatures can be of prognostic and therapeutic value. However, because most of these cancer-related EMT studies are based on bulk gene expression data from clinical specimens, it is often unclear whether clinical EMT signatures originate from mesenchymal malignant cells as opposed to tumor stromal cells (e.g. fibroblasts) that express EMT canonical markers. Furthermore, malignant cells with a purely mesenchymal phenotype appear to be a rare occurrence with a small chance of clinical observation^10^, thus, the existence of such malignant cells is debated.

Adding to the complexity in understanding the clinical significance of EMT is the recognition that EMT is not a binary process (strictly defined by epithelial and mesenchymal states), but instead a continuum of states where transitioning cells exhibit partial EMT phenotypes with both epithelial and mesenchymal features. Partial EMT phenotypes have been observed in clinical cancer specimens and negatively correlate with survival^9,11^, but are poorly understood. Several recent studies have attempted to better define EMT states using single-cell approaches, however they were primarily focused on preclinical models or clinical samples without bridging the two. Pastushenko et al. demonstrated the existence of partial EMT states in mammary and skin cancer by examining a large number of surface markers with flow cytometry and sc-RNAseq^12^. Gonzalez et al. identified partial EMT states in clinical ovarian cancer samples with mass cytometry^13^. While these studies provide important insights, they did not directly relate their findings to EMT states that have been well characterized in cell lines or preclinical reference *in vitro* models. On the other hand, mass cytometry was used to study drug perturbations on TGFβ-induced EMT in mouse epithelial cancer cells^14^ and sc-RNAseq was used to study TGFβ-induced EMT in human breast epithelial cells^15^, but neither of these studies provided a means to assess the clinical relevance of their respective findings.

In this study, we use high-dimensional single-cell analysis, to identify and characterize EMT states observed in primary lung cancer clinical specimens in terms of the spectrum of states observed in well-established lung cancer cell lines. Specifically, through mass cytometry time-course analysis of TGFβ-modulated EMT and MET in lung cancer cell lines, we identify a continuum of EMT and MET states with which we create an EMT-MET STAte MaP (STAMP). In addition, we develop and apply a novel computational trajectory analysis, the TRAjectorty of CElls Reconstruction (TRACER) algorithm, that compares state transitions in EMT and MET. Finally, we develop and apply a machine learning algorithm to project malignant cells from clinical samples onto the EMT-MET STAMP in order to assess clinically relevant EMT and MET states in terms of our *in vitro* time-course analysis. Use of the EMT-MET STAMP, informed by our trajectory analysis, provides a new means to distinguish whether cells are undergoing EMT or MET and thereby has the potential to identify new phenotypic tumor properties, and inform treatment strategies, given that cells undergoing EMT or MET have been shown to exert differential drug sensitivities^9,16^.

## RESULTS

### Phenotyping and ordering epithelial, partial and mesenchymal canonical EMT states in lung cancer cell lines through TGFβ single-cell time-course analysis

To study the spectrum of EMT states in lung cancer cells, we used TGFβ for EMT induction. First, we examined TGFβ responsiveness of 3 NSCLC cell lines (A549, H3255, HCC827) towards bulk expression of commonly used EMT markers ^4^ (E-Cadherin, Vimentin, CD44) (Fig. 1A). A549 cells expressed high levels of both epithelial (E-Cadherin) and mesenchymal (Vimentin, CD44) markers before TGFβ treatment, thereby displaying partial EMT (pEMT) characteristics under basal conditions. With treatment, A549 cells exhibited loss of E-Cadherin and acquired morphological changes associated with a more mesenchymal phenotype (Fig. 1A, Supplementary Fig.1). H3255 cells exhibited primarily epithelial features at basal conditions and largely retained their epithelial morphology and protein expression with TGFβ treatment without displaying EMT changes (Fig. 1A, Supplementary Fig.1). The HCC827 cell line displayed strong epithelial features at basal conditions and underwent dramatic EMT marker and morphology changes with treatment (Fig. 1A-B), hence was chosen as the exemplary cell line for our EMT time-course studies.

**Figure 1.**
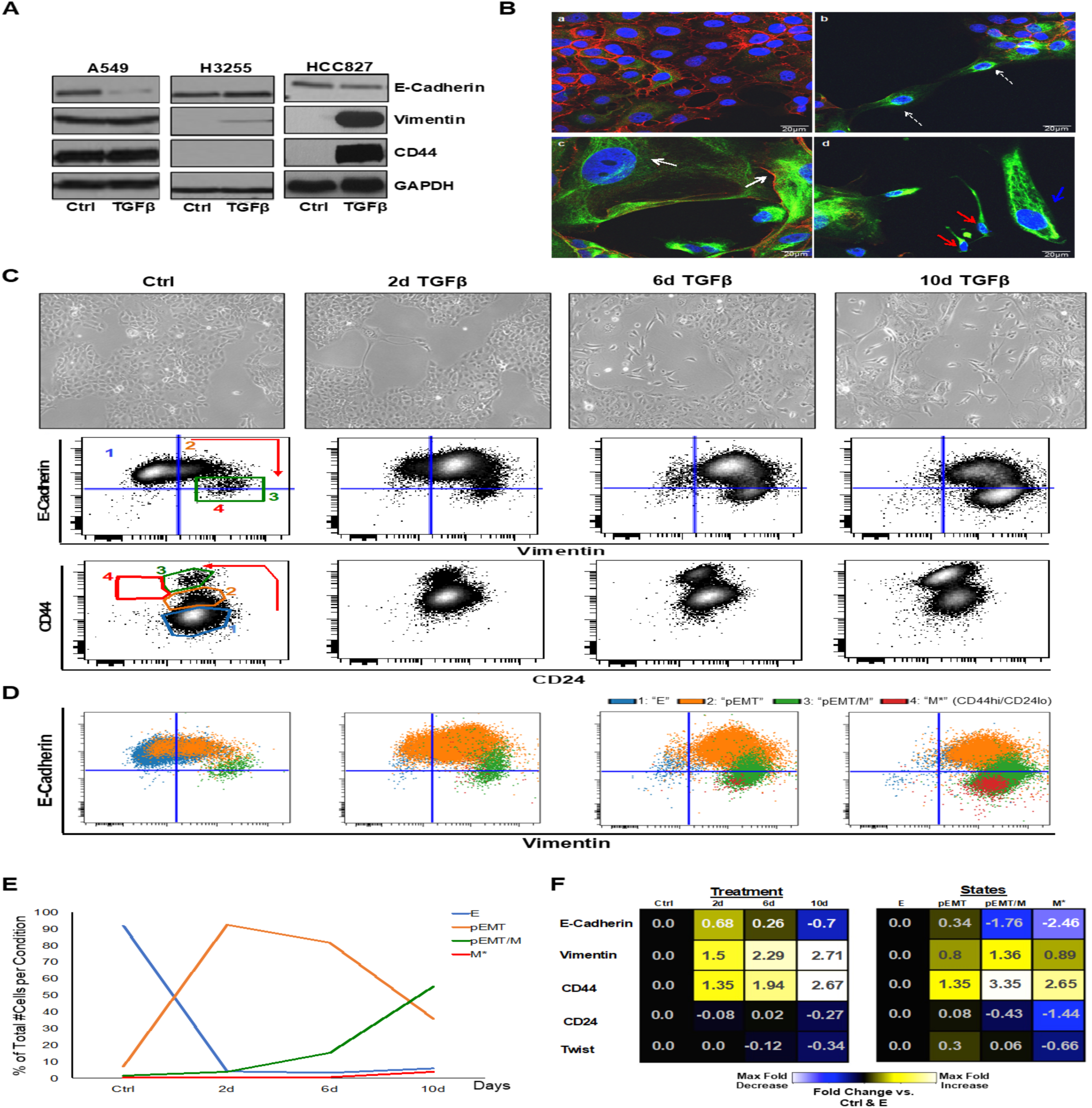
Phenotyping and ordering epithelial, partial and mesenchymal canonical EMT states in lung cancer cell lines through TGFβ single-cell time-course analysis. **(A)** Immunoblots of EMT markers in presence or absence of TGFβ in 3 NSCLC cell lines (1wk treatment in A549 cells, 2wk treatment in H3255 and HCC827 cells). **(B)** Representative confocal images of HCC827 cells stained for E-Cadherin (red) and Vimentin (green) in absence (a, b) or presence of TGFβ (c, d) for 10 days. Magnification 40x. White dotted arrows indicate cells that show loss of E-Cadherin and gain of Vimentin expression in untreated conditions. White solid arrows show cells that have acquired a partial EMT phenotype; blue and red solid arrows indicate cells that have acquired mesenchymal phenotypes. **(C)** Representative images of HCC827 cells treated with TGFβ (5ng/mL) for 2, 6 and 10 days under continuous re-seeding conditions (see also Supplementary Fig. 1 and Methods). Below each image are shown respective E-Cadherin/Vimentin and CD44/CD24 flow cytometry plots. Arrows indicate changes in marker expression during EMT and numbers 1-4 designate the identified canonical EMT states respectively: Epithelial (E), pEMT, pEMT/Mesenchymal (M) and M* (characterized by CD44^hi^/CD24^lo^ CSC-like cells). **(D)** Color-coded gated EMT canonical states depicted on E-Cadherin/Vimentin flow cytometry plots per experimental condition (see also Supplementary Fig. 1). **(E)** EMT state dynamics during TGFβ time-course. Color-coded EMT states were calculated and depicted as % of total number of cells at each experimental day time point. **(F)** Heatmap summary of EMT marker fold changes relative to control condition when analyzed in all cells (left) and relative to the E state when analyzed in the 4 canonical EMT states individually at the 10-day TGFβ time point (right).

Morphological heterogeneity was observed in HCC827 cells during EMT, as evidenced by confocal imaging (Fig.1B). pEMT phenotypes (E-Cadherin^+^, Vimentin^+^) were readily observed in enlarged cells on the periphery of epithelial islands (Fig.1B (c)). Mesenchymal (M) cells (E-Cadherin^−^, Vimentin^+^) appeared either enlarged and elongated or small and spindle-shaped (Fig. 1B (d), blue and red arrows respectively). Intriguingly, cells that were not incorporated in epithelial islands expressed Vimentin even in the absence of exogenous TGFβ, demonstrating that cell density likely affects EMT status (Fig.1B (b)). This finding was corroborated when we examined EMT in HCC827 cells with flow cytometry under varying seeding conditions that controlled confluency (Supplementary Fig. 1). This is consistent with previous reports for normal epithelial cells^17^, but extended here for lung cancer cells. In particular, in sub-confluent conditions, a significantly larger number of cells expressed high Vimentin, of which a subpopulation showed decreased E-Cadherin levels. Transition was also reflected by gradual changes in CD44 and CD24, patterning 4 readily observed EMT states (Supplementary Fig. 1), characterized by progressive increase of CD44. CD24 decreased in only a small subset of cells (CD44^hi^/CD24^lo^), a phenotypic state which has been proposed to be cancer-stem-cell-like (CSC-like)^4,18^.

To adequately capture EMT states, we optimized a TGFβ time-course during which we kept confluency consistent by re-seeding cells at each experimental time point. Cells efficiently underwent EMT within 10 days of TGFβ treatment (Fig.1C, top images). Morphological transition was accompanied by changes in E-Cadherin, Vimentin, CD44 and CD24 expression (Fig. 1C, biaxial plots). Given that these markers are commonly used for EMT characterization, we named the 4 states, canonical EMT states. CD44/CD24 expression in the canonical states 1 through 4 (Fig. 1C, bottom plots) reflected, in terms of E-Cadherin/Vimentin expression the following states: (i) an epithelial (E) (E-Cadherin^+^/Vimentin^−^) state, (ii) a pEMT (E-Cadherin^+^/Vimentin^+^) state, (iii) a mixed pEMT/M (E-Cadherin^+^/Vimentin^+^, E-Cadherin^−^/Vimentin^+^) state, and (iv) a small CSC-like state (E-Cadherin^−^/Vimentin^+^/CD44^hi^/CD24^lo^) denoted here as M* (Fig. 1D, Supplementary Fig. 1). As expected, the E state was most abundant at basal conditions, and rapidly declined by day 2 of TGFβ treatment. Reduction in the E state was accompanied by a sharp increase of the pEMT state at day 2, while pEMT/M and M* states emerged at day 6, with highest numbers at day 10 (Fig. 1E). Mean fold expression of the 4 markers recapitulated the observed changes towards both control conditions with time and towards the E state when examined in each state separately (Fig. 2F). E-Cadherin showed an initial increase during EMT before dropping to its lowest levels at day 10 and this was reflective of E-Cadherin levels in the pEMT state specifically. We speculate that this increase may be the result of increased surface interactions between cells of enlarged size observed during the initial stages of EMT (Supplementary Fig. 1). Unexpectedly, we observed time-dependent changes in Twist, a known EMT transcription factor^19^ (Fig.1F, Supplementary Fig. 1). Specifically, Twist expression was highest in the pEMT state and lowest in the M* state (Fig. 2D, right).

**Figure 2.**
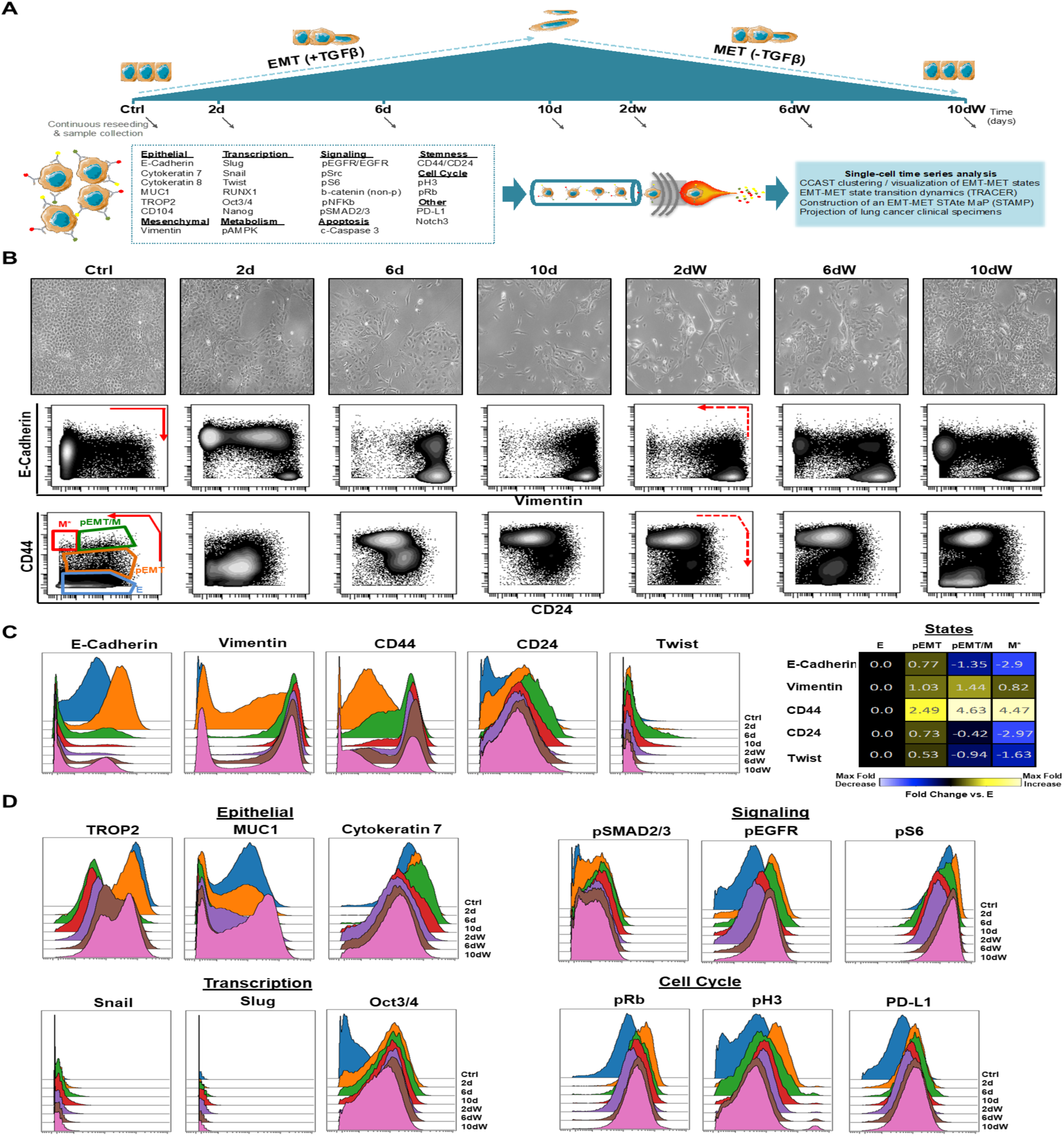
Deep single-cell phenotyping of TGFβ treatment and withdrawal captures composition and timing of canonical EMT states in lung cancer cells. **(A)** Workflow schematic of cell culture conditions for TGFβ time-course (addition and withdrawal (W) time points) and mass cytometry analysis. Arrows indicate times at which cells were collected for both analysis and re-seeding for the remaining time points (see also Supplementary Table 1, Fig. 2–3 and Methods). **(B)** Representative images of HCC827 cells at each time point with respective E-Cadherin/Vimentin and CD44/CD24 mass cytometry plots shown below. Arrows indicate the previously observed canonical EMT marker and states respectively. **(C)** Histogram overlays illustrating EMT marker expression distributions in HCC827 cells undergoing EMT and MET. On the right, a heatmap summary of the same EMT marker fold (arsinh) changes relative to state E when analyzed in the 4 EMT states individually at the 10-day TGFβ time point. **(D)** Additional marker distributions in HCC827 cells undergoing EMT and MET (see Supplementary Fig. 2 for remaining markers and independent biological replicate experiment).

### Mass cytometry analysis provides deep single-cell phenotyping of TGFβ treatment and withdrawal by capturing composition and timing of canonical EMT states in lung cancer cells

For a deeper characterization of the canonical EMT states described above, we performed a mass cytometry time-course experiment on HCC827 cells where TGFβ was added for 10 days and subsequently withdrawn to observe EMT and MET respectively (Schematic Fig. 2A). For optimal confluency conditions, at each time point, cells were collected for both mass cytometry analysis and re-seeding for subsequent time points. In total, 28 markers were measured per cell and were chosen based on literature-derived markers to characterize EMT states and markers that characterize proliferative, signaling and apoptotic cell status (Fig. 2A, Supplementary Table 1). We established consistency between flow and mass cytometry by confirming that the 4 canonical EMT states were reproduced on mass cytometry (Fig.2B, Supplementary Fig. 2). Mass cytometry also reproduced fold expression trends of all previously discussed markers (E-Cadherin, Vimentin, CD44, CD24, Twist) in the identified pEMT, pEMT/M and M* states when compared to E state (Fig. 2C, Supplementary Fig. 2). Upon TGFβ withdrawal, a proportion of cells that had undergone EMT returned to an epithelial state, an observation supported by canonical EMT marker expression and morphological assessment (Fig. 2B-C).

Mass cytometry analysis further enabled us to examine changes of additional markers alongside the canonical EMT markers (Fig. 2D, Supplementary Fig. 2). Keratins and other epithelial cell surface markers (e.g. MUC1, TROP2) showed expected decreases during EMT and subsequent return to basal levels following MET. Most signaling and proliferative markers rapidly increased in expression at day 2 under TGFβ treatment, and subsequently decreased by day 10 in TGFβ. Decreased expression of these markers was sustained during the first 2 days of withdrawal, time points for which M and M* cells were at highest numbers. Specifically, this pattern was observed for pSMAD2/3, a key TGFβ signaling molecule^20^, as well as for pEGFR and pS6, suggesting that signaling is downregulated in the M state. When we examined pEGFR and pS6 levels separately in the 4 canonical EMT states, we found that the M* state was negative for both (Supplementary Fig. 2). This observation is consistent with studies that have associated EMT with tyrosine kinase inhibitor (TKI) resistance^8^ and low p-mTOR activity with stem cell maintenance^21^. We also observed an increase of PDL-1 levels during EMT, substantiating reports that link EMT and tumor immune evasion^22^. Among the measured EMT transcription factors, apart from the transient activation of Twist, we did not detect significant expression levels of Snail or Slug. However, Oct3/4 and Nanog transcription factors showed a substantial increase in expression throughout the EMT time-course; this observation is consistent with a lung adenocarcinoma study where co-expression of Oct3/4 and Nanog were found to be critical for EMT^23^.

### Unsupervised analysis of high-dimensional mass cytometry data reveals new computationally-derived EMT states and identifies a previously unrealized partial MET state in lung cancer cells

Given the high dimensionality of our mass cytometry EMT-MET time-course data, we applied unsupervised cluster analysis to computationally derive a continuum of EMT and MET states. From hereon, the canonical EMT states (E, pEMT, pEMT/M, and M*) hold no particular significance but will be related to computationally-derived EMT states. We first applied a clustering algorithm, CCAST, on the pooled mass cytometry data across all time points^24^. E-Cadherin, Vimentin, CD44, CD24, MUC1 and Twist were selected to identify the clusters based on PCA (Supplementary Table 2). CCAST identified 8 most prominent states, with each state having ≥1% of total number of cells analyzed. CCAST and downstream analyses, as well as the relation between the computationally-derived states with the 4 canonical EMT states are shown in Supplementary Fig. 3-4.

Heatmaps of the 6 clustering markers within each of the 8 states revealed minimal within-state marker heterogeneity compared to between-state marker heterogeneity (Fig.3A). We labeled the derived EMT and MET states based on their expression profile as well as their respective time-course pattern. Among the 8 states (denoted here in italics), we identified 3 epithelial states (*E1, E2, E3*), 3 partial EMT states (*pEMT1, pEMT2, pEMT3)*, one mesenchymal state (*M*) and one partial MET state (*pMET*) (Fig. 3B). The **epithelial states** *E1, E2* and *E3* were consistent with canonical epithelial marker expression, where all displayed low expression of Vimentin and CD44 but revealed varying degrees of E-Cadherin and CD24 expression. MUC1 was high in *E1* and *E2* and absent in *E3*, similar to all other markers in *E3* except E-Cadherin. All 3 epithelial states were highest in numbers at time 0 and dramatically decreased with TGFβ treatment; only *E1* and *E2* rebounded following TGFβ withdrawal. The **partial EMT states** *pEMT1, pEMT2*, and *pEMT3* were all positive for E-Cadherin and Vimentin, consistent with a pEMT phenotype. However, compared to *pEMT1, pEMT2* and *pEMT3* showed increased levels of Vimentin and CD44, with *pEMT3* also showing a dramatic decrease of the epithelial marker MUC1. Furthermore, *pEMT2* and *pEMT3* each included a subgroup of Twist^+^ cells (Fig. 3A boxed regions). Twist^+^ cells in *pEMT3* expressed lower E-Cadherin and MUC1 levels than Twist^+^ cells in *pEMT2*, implicating an EMT switch within Twist^+^ cells specifically during treatment. All partial EMT states were absent at time 0 and emerged upon TGFβ treatment, decreased with continued TGFβ treatment, and with the exception of *pEMT3*, transiently re-emerged during TGFβ withdrawal. The **mesenchymal state** *M* was characterized by loss of epithelial markers (E-Cadherin, MUC1), and increased expression of both Vimentin and CD44, and was highest in numbers at day 10 in TGFβ. Interestingly, no Twist^+^ cells were detected within this state, confirming a study that found that CSC-like properties arise following transient Twist activation during EMT^25^. Although our clustering analysis did not distinguish CSC-like cells as a separate cluster, we did observe heterogeneity among *M* cells and interrogation of other markers in cells expressing the lowest CD24 levels (e.g. pS6) helped us define the CSC-like cells within the *M* state (*M\**, Supplementary Fig. 4). The **partial MET** state *pMET*, perhaps most novel in our analysis, is a previously uncharacterized state that peaked during withdrawal time points, representing cells that had perhaps initiated MET. The *pMET* state displayed a similar phenotype with the *M* state in 5 of the 6 markers but was significantly higher in MUC1 expression.

**Figure 3.**
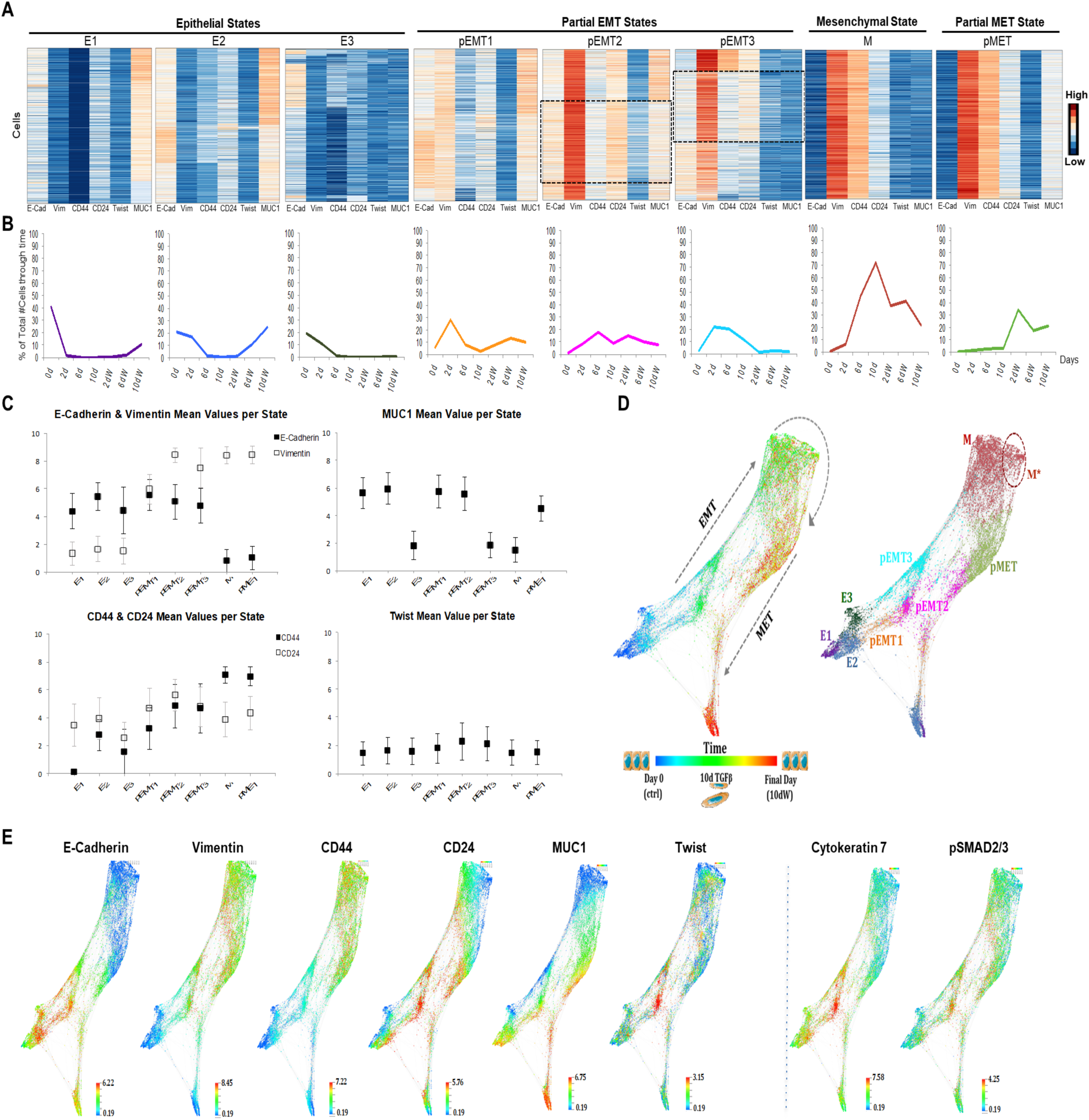
Unsupervised analysis of high-dimensional mass cytometry data reveals new computationally-derived EMT states and a unique partial MET state in lung cancer cells. **(A)** Heatmaps depicting the expression of the 6 clustering markers in each cell (E-Cadherin, Vimentin, CD44, CD24, Twist and MUC1) per computationally-derived EMT and MET state (normalized global expression). Shown are the most prominent in numbers states that resulted from the CCAST algorithm performed on pooled mass cytometry HCC827 data. Dotted boxed regions illustrate subpopulations of Twist^+^ pEMT cells. (See also Supplementary Fig. 4). **(B)** EMT and MET state dynamics. Each graph corresponds to the state heatmap above. States were calculated and depicted as % of total number of cells at each experimental time point. **(C)** Graphs illustrating mean expression of the 6 clustering markers in EMT and MET states. Bars represent standard deviation of each marker within each state. **(D)** Time-resolved force-directed (FDL) layout of mass cytometry time-course data (left) and respective EMT and MET state annotations (right). Dotted area depicts subpopulation of M cells that exhibit CSC-like phenotypic characteristics (CD44^hi^/CD24^lo^, *M\**). **(E)** Time-resolved FDL layouts colored by protein expression of indicated markers (Arsinh transformed data). See Supplementary Fig. 4 for remaining markers and analysis of an independent biological replicate experiment.

To interrogate the temporal dynamics of the computationally derived EMT and MET states, we pursued a variety of approaches that quantify changes over time. First, we ordered the states according to their temporal pattern and phenotype and quantitatively examined the expression changes of the 6 clustering markers (Fig. 3C). Vimentin levels increased sharply upon the appearance of the pEMT states (*pEMT1, pEMT2, pEMT3)*, preceding the drop of E-Cadherin. CD44 gradually increased reaching its highest levels in the *M* and *pMET* states. CD24 expression varied in the various states, with lowest levels observed in states *E3* and *M*. Confirming our initial observations, Twist expression peaked in the states *pEMT2* and *pEMT3*, followed by a drop in states *M* and *pMET*, comparable to levels observed in epithelial states. MUC1 displayed dramatic expression differences, with lowest levels observed in states *E3, pEMT3* and *M* and high levels in states *E1, E2, pEMT1, pEMT2* and *pMET*. Similar results were observed in an independent biological replicate experiment (Supplementary Fig. 4).

To better visualize the phenotypic profiles of the derived EMT and MET states and their dynamic changes, we constructed a force-directed layout (FDL) of the high-dimensional data using Vortex^26^ (Fig. 3D, Supplementary Fig. 3). Cells from each of the 8 computationally-derived states were added sequentially per time point and data were visualized in terms of time (Fig. 3D, left panel), or state (Fig. 3D, right panel). Thus, we were able to visually decipher the EMT from the MET trajectory, and identify states which appeared to be transient (e.g. *pEMT3*), or revisited during MET (e.g. *pEMT1, pEMT2*), or at MET completion (e.g. *E1, E2*). Strikingly, *pMET* state appeared to be specific to MET, supporting studies proposing that MET differs from EMT^27,28^. The expression profiles of all markers are depicted in FDLs in Fig. 3E and Supplementary Fig. 4. FDLs reveal that Twist^+^ cells are characterized by high expression of cytokeratins 7 and 8, signaling molecules (e.g. pSMAD2/3, pEGFR), and EMT-related transcription factors Oct3/4 and Nanog. High cytokeratin expression corroborates studies showing that Twist^+^ cells constitute a specific type of epithelial cells^29^. On the other hand, cells in the *M* state are characterized by low signaling and proliferative profiles, a finding that agrees with our earlier observations (Fig. 2, Supplementary Fig. 2) and published reports^21,30^.

### Construction of an EMT-MET STAte MaP (STAMP) and TRAjectory of CElls Recontruction (TRACER) analysis identifies distinct EMT and MET trajectories and state transition dynamics in lung cancer cells

Apart from visualizing the phenotypic, time-dependent properties of EMT and MET states, our ultimate goal was to construct a EMT-MET STAte MaP (STAMP) for inferring the presence of our 8 derived EMT and MET states in lung cancer clinical specimens with single-cell resolution. To construct the EMT-MET STAMP, we first generated a 2D t-SNE projection of our derived EMT and MET states from our TGFβ mass cytometry study. Next, to define regions of the map most uniquely associated with each state, we determined the highest density region per state per sampling time and estimated its respective centroid. Given that transition is a continuum of EMT states with considerable overlap, we employed Voronoi and Convex Hull analysis^31^ to achieve density-driven segmentation of the EMT t-SNE landscape; each segment corresponding to an identified EMT/MET phenotypic state at its peak during our time-course analysis (Fig. 4A-B and Supplementary Fig. 3, 5). Analysis of cell density over time on the EMT-MET STAMP confirmed our previous assessment that MET differs from EMT and is specifically characterized by the appearance of the *pMET* state at the 2-day withdrawal time point (Fig. 4C). However, it also revealed the possibility that cells that belong to states *pEMT2* and *pEMT3* exhibit plasticity that enables them to follow the same state trajectory back toward *pEMT1* and *E* states under TGFβ withdrawal. This observation is heuristically demonstrated in a density-driven assessment of transitions within the 2D map of EMT-MET states, depicted in Fig. 4D. When we applied the pseudotime/cell lineage computational tool Slingshot^32^ to this data, it independently resolved the trajectory that involves the appearance of the *pMET* state during withdrawal (Supplementary Fig. 5), however, it did not fully capture our empirical observations of likely state transitions.

**Figure 4.**
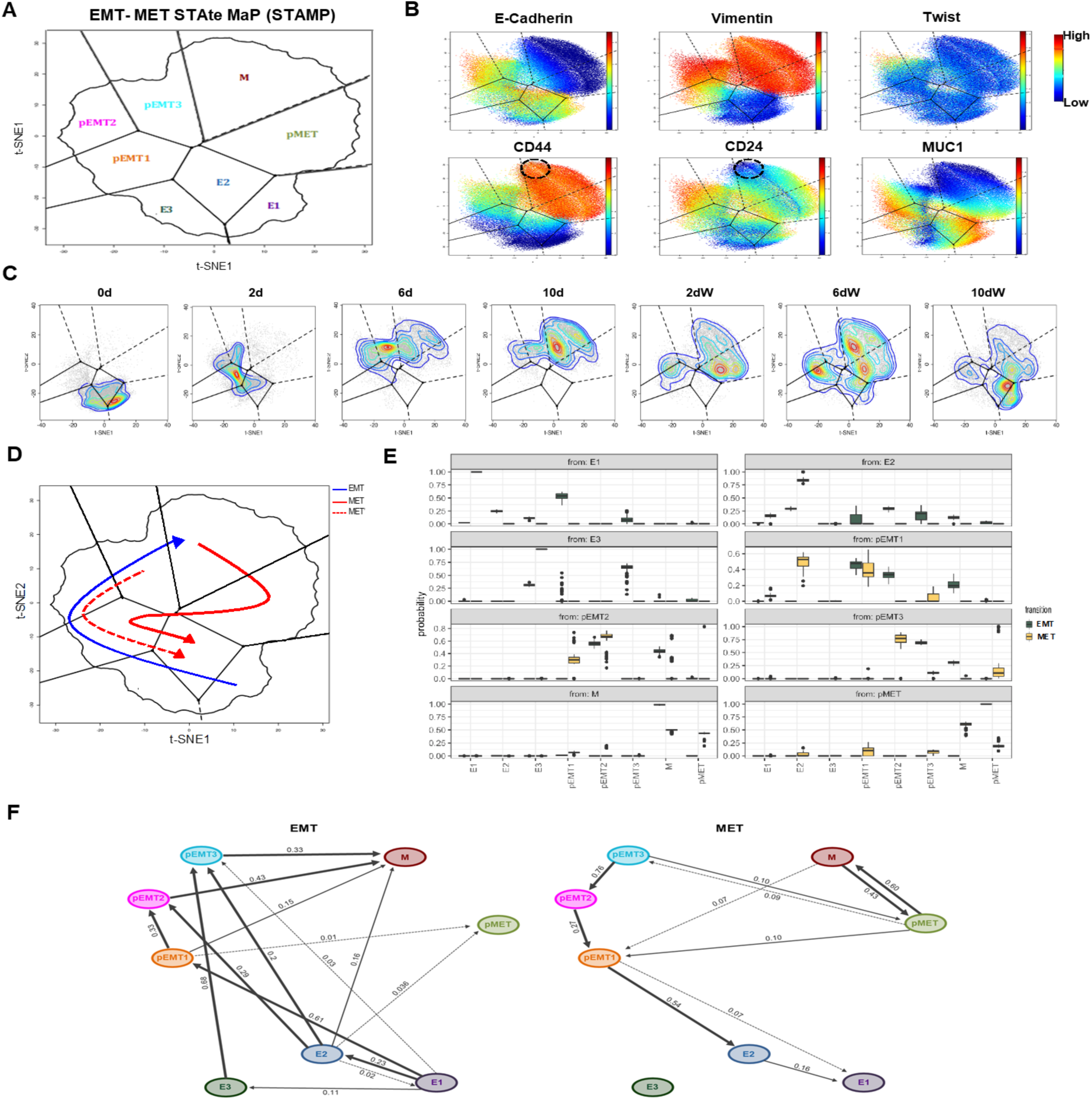
Construction of an EMT-MET STAte MaP (STAMP) and TRAjectory of CElls Recontruction (TRACER) analysis identifies distinct EMT and MET trajectories and state transition dynamics in lung cancer cells. **(A)** Schematic of 2D t-SNE EMT-MET STAte MaP (STAMP). HCC827 CCAST state information was used to determine the highest-density areas of cluster-specific time dependent bins and subsequent estimation of respective centers. Compartmentalization was achieved with Voronoi and Convex hull analysis. **(B)** Expression profiles of the 6 clustering markers in pooled HCC827 time point data visualized on the EMT-MET STAMP. See Supplementary Fig. 5 for remaining markers. **(C)** Time point specific HCC827 t-SNE density plots. **(D)** Schematic hypothetical model of possible EMT and MET trajectories of transitioning cells (see text for details). Blue arrow depicts the EMT trajectory, whereas red arrows depict 2 possible MET trajectories that both may take place depending on conditions; one utilizes a new state (*pMET*) which supports the notion of hysteresis during MET (red solid line), the other utilizes the previously visited *pEMT* states (red dotted line). **(E)** Bootstrap analysis comparing the distribution of time independent state transitions as generated by the TRAjectory of CElls Recontruction (TRACER) algorithm for EMT and MET, represented as box plots, each graph showing transition of a specific state to the all other states (x axis). The centers of the box plots represent medians of the bootstrap transition distributions. Box plots color coded as green (left) and yellow (right) represent EMT and MET, respectively. **(F)** Schematic diagram of the TRACER medoid network from bootstrap analysis, depicting transitional probabilities among states during EMT and MET. Arrows are weighted by probability strength. See Methods for further details.

To more rigorously assess the potential state transitions, we developed a novel computational trajectory reconstruction algorithm, TRACER (Supplementary Fig. 3 and Methods) that does not rely on pseudotime assumptions and allows branching. We assumed that EMT and MET are classical Markov processes^33^ with transition probabilities that are constant between the states within EMT and MET but can differ when comparing EMT and MET. However, because our observations of the proportion of cells in each state are made at discrete, non-uniform time points, and because we do not observe the trajectories of individual cells, we cannot employ standard methods to estimate for the state transition probabilities^34^. Also, because the number of states is relatively large compared to the number of observations, we employ sparsity assumptions^35^. We modeled the state transition probabilities *p*_*jkt*_ between states *j* and *k* at time *t* by imposing sparsity penalties on *p*_*jk*t_ for *j* not equal to *k*, but no penalty on *p*_*jjt*_, with the idea that there is no cost for staying in a state, but switching between states is discouraged. Under these assumptions, EMT and MET state transition probabilities can be estimated by convex optimization, with the sparsity parameter λ selected through cross-validation. We applied our model separately for the EMT and MET time points and, through bootstrap analysis, we generated a distribution of the transition probabilities between states under EMT and MET, as shown in Fig. 4E. In general, the bootstrap analysis shows less non-zero state transition probabilities under MET compared to EMT, clearly demonstrating that the transition between *M* and *pMET* states is unique to MET. The medoid networks for EMT and MET are provided in Fig. 4F. Interestingly, during MET, all cells transitioning from either a *pEMT* or the *pMET* state are estimated to transition through the *pEMT1* state before becoming epithelial. These results demonstrate statistically significant differences between EMT and MET trajectories as well as bidirectionality (plasticity) between certain states (e.g. between states pEMT1 and pEMT2).

### Projection onto EMT-MET STAMP using a neural net algorithm serves as a means to assess EMT-MET states in independent samples and is verified using NSCLC cell lines

Given the wide range of phenotypic, and presumably functional, EMT-MET states across NSCLC cell lines, we tested whether our EMT-MET state map could serve as a reference to reconcile known heterogeneity between cell lines and experiments. To project independent samples onto the EMT-MET STAMP, we trained a neural net algorithm^36,37^ that predicts the bivariate t-SNE outputs (i.e. tSNE1 and tSNE2) of our EMT-MET STAMP in terms of the selected 6 clustering markers (Supplementary Fig. 3 and Methods). We verified the performance of our neural net mapping function using lung cancer cell lines, which have known features. First, we projected independently analyzed TGFβ HCC827 samples, and these matched previously observed mappings (Fig. 5A, left). Next, we projected two independent NSCLC cell lines (A549, H3255), given that we had prior knowledge of their EMT status (Fig.1A). As expected, A549 cells mapped onto the *pEMT3* and *M* region in control conditions and extended into the *M* region upon TGFβ treatment (Fig. 5A, middle). On the other hand, and in agreement with our previous findings, H3255 cells mapped mostly on *E2*, and to a lesser degree on *pEMT1*, with almost no changes upon TGFβ treatment (Fig. 5A, right). These map projections were consistent with morphological assessment and respective E-Cadherin/Vimentin and CD44/CD24 expression profiles (Fig. 5B-C).

**Figure 5.**
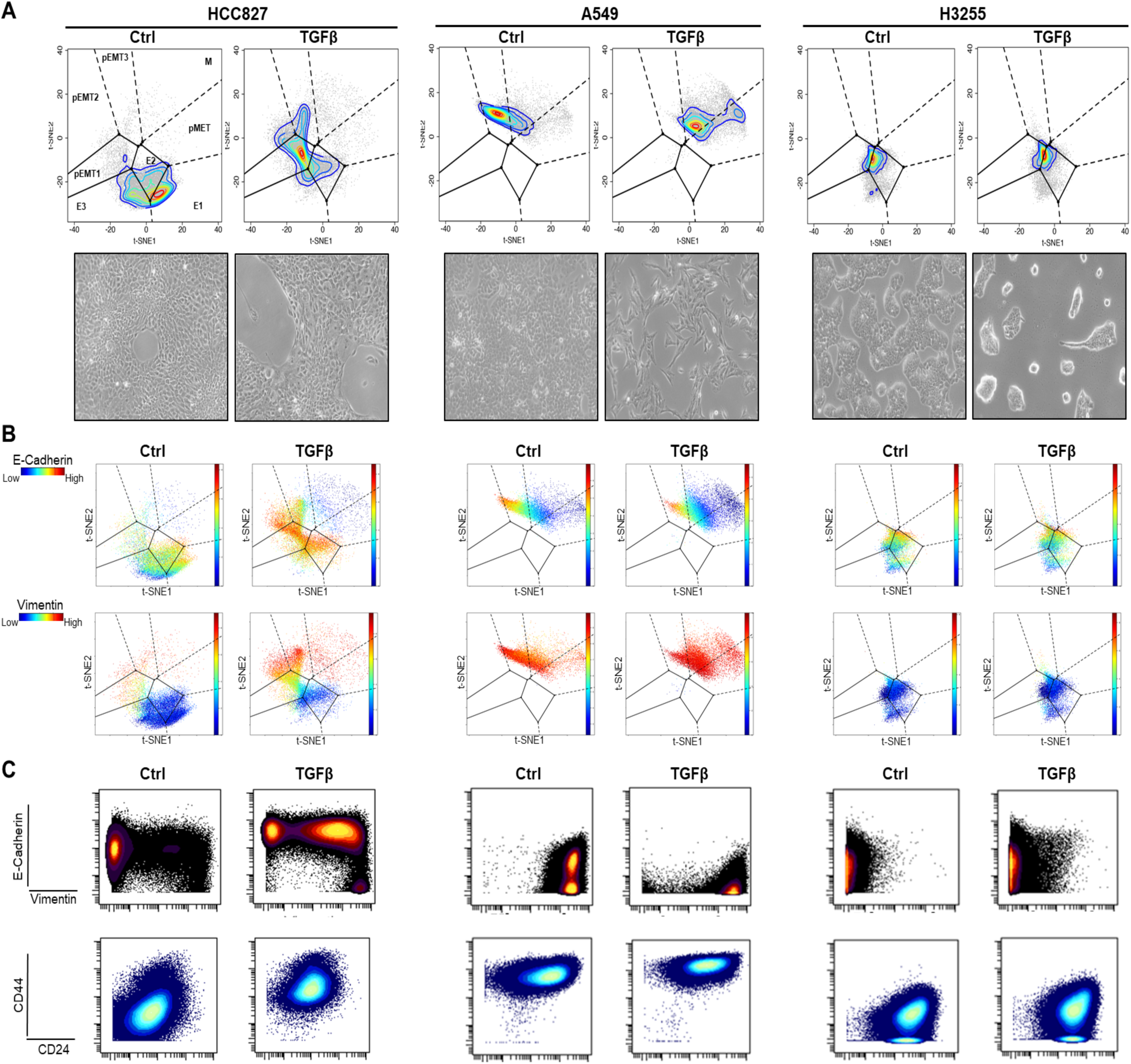
Projection onto EMT-MET STAMP using a neural net algorithm serves as a means to assess EMT-MET states in independent samples and is verified using NSCLC cell lines. **(A)** Projection of NSCLC cell lines onto the EMT-MET state map (top) and their respective morphological assessment (10x images, bottom). HCC827 samples (ctrl, 2-day TGFβ) were ran on a different day than the ones used for the map construction for validation (left). A549 and H3255 cells were treated with TGFβ for 10 and 13 days respectively. **(B)** Shown are the respective to each cell line and condition expression profiles of E-Cadherin (top) and Vimentin (bottom) visualized on the t-SNE EMT-MET STAMP on which they were projected. **(C)** E-Cadherin/Vimentin and CD44/CD24 mass cytometry plots per projected cell line sample. See Methods for further details.

### Projection of clinical NSCLC samples onto the EMT-MET STAMP demonstrates the existence of the partial and full EMT and MET states

We proceeded to determine the spectrum of EMT and MET states in lung cancer clinical specimens with single-cell resolution by projecting them onto the EMT-MET STAMP using our neural net algorithm described above. Five fresh NSCLC adenocarcinoma samples were obtained immediately after patient resection under IRB approval and underwent immediate dissociation for single-cell suspension. For mass cytometry analysis, we used the antibody panel developed for our time-course analysis, augmented with antibodies to sort out immune (CD45^+^), endothelial (CD31^+^) and stromal fibroblast (FAP^+^) populations^13^. With this, we were able to discriminate immune, endothelial and stromal cells from cells that were of varying levels of cytokeratins 7 and 8, while at the same time negative for CD45, CD31 and FAP (Supplementary Fig. 6). The specimens harbored different mutations and their differentiation status was graded as poor, moderate or well in the pathology report. Tumor size and smoking history were also recorded (Fig. 6, Supplementary Table 3). We validated both the mass cytometry run and the projection analysis, by analyzing in parallel independent HCC827 samples as a control (Supplementary Fig. 6). Upon projection onto the EMT-MET STAMP, all clinical specimens showed presence of a variety of EMT states that agreed with their respective E-Cadherin/Vimentin expression profiles (Fig. 6, Supplementary Fig. 6). Reassuringly, given the single-cell resolution of the mapping function, each clinical sample demonstrated a continuous sweep of EMT states, as opposed to occupying disjoint regions. Cases No. 1, 2 and 3 (all EGFR mutated) mapped primarily onto epithelial regions with some extent into the *pEMT1* regions, and expressed, as expected, higher pEGFR levels (Supplementary Fig. 7). Interestingly, this mapping is consistent with EGFR mutated cell lines HCC827 and H3255 in basal conditions and confirmed by morphological assessment (Supplementary Fig. 6). Case No. 3, harboring an additional TP53 mutation, broadly extended into the pEMT regions, primarily occupying the *pEMT3* region, with a small fraction of cells in the *M* and *pMET* regions. Case No. 4 and 5, harboring TP53 and KRAS mutations respectively, showed the most partial EMT features, occupying the *pEMT2* and *pEMT*3 regions (and to a lesser degree *M*), reassuringly similar to A549 cells that are also KRAS mutated (Fig. 5). Interestingly, Case Nos. 4 and 5, in comparison to Case Nos. 1-3, were graded as poorly differentiated and showed higher levels of immune infiltration (Fig. 6, Supplementary Fig. 6), suggesting the association of these clinical features with specific EMT states. While a larger sample size is necessary to solidify conclusions, these results are consistent with studies reporting correlations between NSCLC mutations and immune infiltration with EMT status^22,38^. Overall, we show that our EMT-MET STAMP can be utilized to score and interpret clinical specimen data towards EMT and MET state heterogeneity and that similar approaches can be extended to phenotyping a range of cellular processes that involve cell state transitions in various types of cancers.

**Figure 6.**
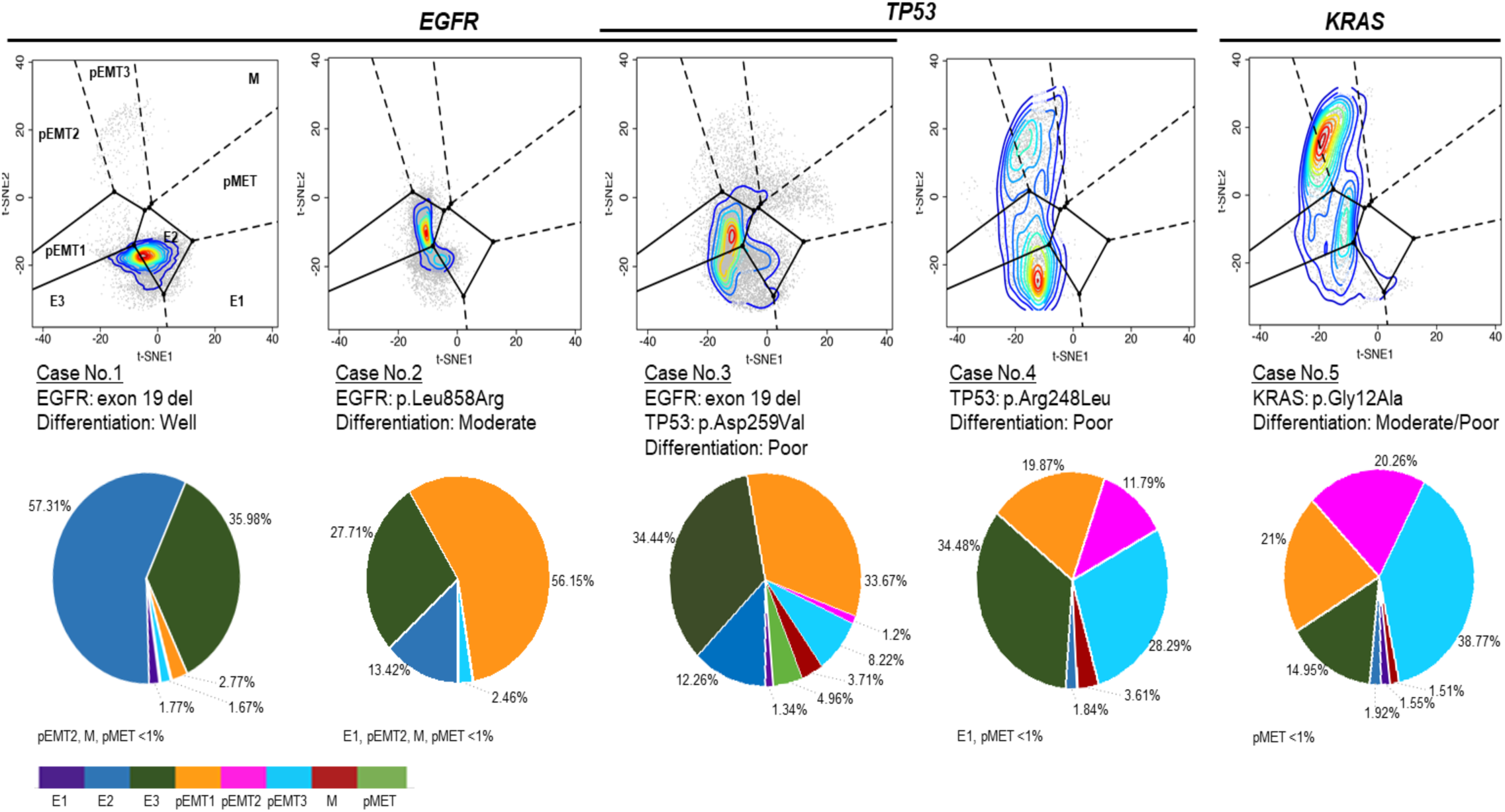
Projection of clinical NSCLC samples onto the EMT-MET STAMP demonstrates the existence of the partial and full EMT and MET states. Shown are the projections of 5 clinical samples analyzed with mass cytometry and their respective mutation and differentiation features. Pie charts show % of tumor cells per clinical sample mapping on the color-coded computationally-derived EMT-MET states. See Methods, Supplementary Table 3 and Fig. 6-7 for further details.

## DISCUSSION

Elucidating a continuum of states in EMT and MET in clinical samples promises new insights in cancer progression and drug response. In order to assess the clinical relevance of EMT and MET properties gained from extensive *in vitro* studies, we provide a novel integrated experimental-computational framework. First, we identified a continuum of EMT and MET states leveraging high-dimensional single-cell mass cytometry time-course analyses of lung cancer cell lines undergoing EMT through TGFβ-treatment and MET through TGFβ-withdrawal. To translate the potential clinical significance of our *in vitro* findings, we constructed an EMT-MET state map (STAMP), and using a neural net algorithm, we projected clinical samples onto the map to evaluate their EMT and MET state profile with single-cell resolution. This integrated experimental-computational approach provides novel *in vitro* insights on EMT-MET biology and establishes a novel framework to translate these findings in clinical samples.

Our mass cytometry time-course analysis of TGFβ-modulated EMT and MET provides novel biological insights. First, we demonstrate not only the heterogeneity of EMT cell states but also transient properties that provide a deeper appreciation of the dynamism of the EMT process. For example, we showed that in lung adenocarcinoma epithelial states (i.e. *E1, E2, E3*) can be quite heterogeneous, displaying varying levels of E-Cadherin, CD24 and, interestingly, MUC1. Heterogeneity was also observed within the partial EMT states for various markers, most notably Twist, with a partial EMT state (e.g. *pEMT1*) emerging prior to the increase of Twist expression. Expression of EMT-specific transcription factors during EMT was recently reported to be dispensable for some cells, featuring an alternative EMT program that involved protein internalization^40^. We observed phenotypically distinct but transient Twist^+^ cells that retained some epithelial identity and proliferative capacity, consistent with prior work^29^. Although transient Twist activity has been reported^25^, these prior observations were made in cells where Twist was exogenously overexpressed; instead, we examined protein changes in *endogenous* Twist under a well-accepted mode of EMT induction (TGFβ). Our finding is clinically provocative because Twist has been shown to be overexpressed in human lung adenocarcinoma and specifically correlated to EGFR mutations^41^, as observed in two of three EGFR mutated clinical samples we analyzed (Supplementary Fig. 6). Even though we did not detect expression of other EMT-specific transcription factors (i.e., Slug, Snail, Zeb1, Supplementary Fig. 1), we cannot exclude the possibility that these may be activated in earlier EMT time points not tested here.

Our time-course analysis of MET is a key aspect of our study. MET is thought to be critical to the establishment of secondary distant tumors. Yet, compared to EMT, MET is less studied, particularly with single-cell resolution. Some studies have shown that EMT is reversible among cells in partial EMT states but not necessarily among cells that have become mesenchymal, although this seems to be cell-type dependent^42,43^. Even so, for cells undergoing MET, it is unclear whether the MET trajectory mirrors or differs from the EMT trajectory. Differing trajectories that we found is evidence of hysteresis, a phenomenon in which the dependence of a future state depends on its history. Several mathematical modeling studies have provided evidence of hysteresis when comparing EMT and MET, however these were based on gene expression data or were not associated with specific phenotypic states^27,28^. By analyzing mass cytometry time-course data using our novel algorithm TRACER, we find statistically significant evidence of hysteresis. In particular, we showed that some mesenchymal cells undergo MET utilizing a trajectory not observed under EMT and transit through a newly identified state that we defined as *pMET*. More specifically, our study supports two possible scenarios. In the first scenario, cells in the M state have undergone such significant (presumably epigenetic) changes that in order to undergo MET, they utilize a different trajectory. Consistent with this, is the likelihood that M cells are more stable than partial EMT cells, given that a significant proportion of cells failed to undergo MET after TGFβ withdrawal. Moreover, we found that if cells have not efficiently undergone EMT (majority of cells transition to pEMT and not M states), most of them are able to undergo MET (Supplementary Fig. 5). In the second scenario, if during conditions that promote MET a cell is in a partial EMT state, it utilizes a mirrored trajectory back to an epithelial state. Supporting this, the medoid network from TRACER, and the bootstrap analysis, detected bidirectionality between the partial EMT states (Fig.4E-F). Future studies involving live cell tracing and Twist knockout experiments will be critical towards deciphering which exact populations undergo complete EMT, retain plasticity and/or undergo hysteresis/MET.

Our study introduces a neural net algorithm for projecting samples on the EMT-MET STAMP with single-cell resolution. The EMT-MET STAMP can be used as a potential tool to assess NSCLC clinical specimens in terms of their EMT status, as defined along a well-established and reproducible *in vitro* time-course analysis. Our mapping of five clinical samples serves as proof of concept that such mapping can offer insights on the clinical relevance of the spectrum of EMT and MET states. For example, we observed a potential association of EMT with immune infiltration. Such a correlation has been previously reported but not linked to mutation status^44^, supporting not only the necessity of a larger scaled study, but the importance of a well-defined EMT reference map. Perhaps the most intriguing aspect of projecting onto the EMT-MET STAMP, is the promise it holds for future therapeutic efforts. EMT and MET have been correlated with drug resistance^8,45,46^. Therefore, being able to assess malignant cells from clinical samples in terms of specific EMT and MET states can be informative. For instance, some studies have suggested that inducing MET could be beneficial by restoring sensitivity to TKI inhibitors^16,47^. Indeed, our data showed that pEGFR signaling was lowest in the *M* (specifically M*) state, whereas in cells undergoing MET, pEGFR levels increased (Supplementary Fig. 2-5), which would hypothetically constitute TKI sensitivity. Therefore, depending on the situation and timing, one may choose to target or induce the *pMET* state, and having a proteomic-based EMT-MET map featuring targetable markers could benefit such efforts. To this end, our map showed that we could theoretically distinguish whether a specimen had cells undergoing EMT or MET, and this alone could have implications on clinical assessment. Specifically, our data show that if cells are in the *pMET* state, they significantly differ from M cells primarily in terms of MUC1 expression, a molecule that is being studied as a therapeutic target in lung cancer^48^. None of our clinical specimens mapped significantly onto the *pMET* region, barring Case No 3. This was expected, because such cells are more likely present in metastatic and not primary tumors; however, this does not negate the possibility that cells within any given tumor may undergo MET depending on spatiotemporal conditions. In addition to predicting drug resistance, an EMT-MET map may have utility for evaluating disease progression and drug response during therapy at the individual patient level. For example, analyzing malignant pleural effusions has been proposed as means to evaluate disease status and EMT markers have been detected in lung cancer pleural effusions^49,50^.

Our study proposes a novel approach for interrogating and mapping EMT states but has its limitations. Although here we chose TGFβ to induce EMT in lung cancer cells, it is well known that EMT can be triggered by a variety of conditions including hypoxia and drug treatments^51^. To expand the concept that we presented, it would be reasonable to test and compare different modes of EMT induction and examine how these would affect the constructed state map, or the projections onto it. However, it has been reported that the EMT phenotypes acquired under different inducers are quite common, barring some signaling pathway differences^51^; suggesting that our map, that was built on canonical EMT markers, could have broader use. Moreover, the fact that common EMT phenotypes exist despite the variety of signaling/mutational underlying backgrounds, justifies our choice to build our map with the HCC827 cell line, as it portrayed a complete continuum of EMT-MET states. Nevertheless, constructing an EMT-MET state map that incorporates different mutational backgrounds and modes of induction is a worthy pursuit. It is important to note that projecting individual specimens on a reference map differs from traditional approaches where multiple samples are pooled for analysis; allowing each specimen to be examined within its own heterogeneous identity. Also, by using single-cell proteomic data, our approach differs from similar studies that have employed basic concepts from machine learning to classify and diagnose clinical samples based on gene profiling alone^36,52^. Finally, our work can be extended to multiplexed imaging^53^ to further delineate the relationships between EMT in cancer cells and their surrounding spatiotemporal/microenvironment elements in unmodified tissue.

In summary, we have defined a landscape of EMT and MET states with single-cell resolution from an *in vitro* time-course analysis where we modulated EMT states with TGFβ treatment and withdrawal. With this information, we created an EMT-MET state map (STAMP) as a new means to assess clinical samples towards prognosis and predictive efforts, specifically when assessing therapeutic response and drug resistance. Furthermore, our experimental and computational approaches provided insights on aspects of EMT basic biology, EMT/MET trajectories and the heterogeneity of transient cell subpopulations undergoing phenotypic transitions. Combining our work with functional characterization of the states we have presented here (e.g. correlating a specific state with drug resistance), and incorporation of mutational and clinical information across more patient samples will enable a more a comprehensive understanding of the diagnostic, prognostic and therapeutic relevance of EMT in cancer.

## ACKNOWLEDGMENTS

We gratefully acknowledge Dr. Wendy J. Fantl for discussions and technical advice on analyzing primary clinical specimens with mass cytometry and for a valuable review of our manuscript. We thank Dr. Matt van der Rijn for discussions and input on NSCLC histology and pathology, Kelsey Ayers for assisting initial clinical specimen acquisition, Dr. Tyler Risom for his input on data presentation and Trevor Bruce for mass cytometry technical assistance. We thank Dr. Parag Mallick for providing the NSCLC cell lines used in this study.

## CONTRIBUTIONS

Conceptualization, L.K.G., B.A., S.C.B., S.K.P.; Methodology, L.K.G., S.C.B., B.A., N.I., R.T., S.K.P.; Investigation, L.K.G., S.C.K.; Software and Formal Analysis, L.G.K., B.A., N.I., R.T., S.K.P.; Resources, J.A.B, J.B.S, S.C.B., S.K.P.; Visualization, L.G.K, B.A., S.K.P.; Writing-Original Draft, L.G.K., S.C.B., S.K.P.; Writing-Review & Editing, All authors; Supervision: S.C.B., S.K.P.; Funding Acquisition, B.A., S.C.B., S.K.P.

## METHODS

### Cell line Studies

HCC827, H3255 and A549 NSCLC adenocarcinoma cell lines were a generous gift from Dr. Parag Mallick and were grown in RPMI and DMEM media respectively, supplemented with 10% fetal bovine serum (FBS) and 5% antibiotic solution (penicillin/streptomycin), at 5% CO2 and 37°C.

### Human Studies

Clinical aspects of this study were approved by the Stanford Institutional Review Board in accordance with the Declaration of Helsinki guidelines for the ethical conduct of research. All patients involved provided informed consent. Collection and use of human tissue was approved and was in compliance with data protection regulations regarding patient confidentiality. Following surgical resection of primary tumors from five patients at Stanford hospital, NSCLC adenocarcinoma specimens were immediately processed in order to achieve single-cell suspensions for mass cytometry analysis.

### EMT Induction in NSCLC cell lines

For optimal EMT induction cells were seeded in 100mm tissue culture plates at 750 000 cells/plate for 24hrs and then switched to 2% FBS media for 24hrs. Cells were then treated with 5ng/mL TGFβ (R&D, #240-B/CF) for various time points in 2% FBS media. To keep cell density a non-critical factor for EMT induction, cells were re-seeded (same number of cells/plate) at each collection time point through the entire course of the treatment. For withdrawal conditions, cells were seeded from TGFβ treated cells in absence of ligand and collected at same time increments as with TGFβ treatments with continuous re-seeding as mentioned above. In certain cases, TGFβ was added to cells for a period of 1 or 2 weeks with or without re-seeding as indicated in respective figure legends.

### Immunoblotting

Cells were lysed by adding lysis buffer (50mM Tris, pH 8.0, 2% SDS, 1x protease inhibitor cocktail, 25mM NaF, 100uM Na_3_VO_4_, 5mM EGTA and mM EDTA) to adherent cells (using a cell scraper). Lysates were then sonicated on ice in order to shear DNA and reduce viscosity. To pellet cellular debris, lysates were centrifuged at 14000 rpm at 4°C for 10 minutes and supernatants were transferred to fresh tubes. Protein quantification was performed using the Pierce BCA protein assay kit (Thermoscientific) and equal amounts of total protein were subjected to standard electrophoresis conditions. Separated protein lysates were transferred to PVDF membranes (Millipore #IPVH00010), which were subsequently rinsed in TBST (Tris-buffered saline, 0.1% Tween 20). Blocking non-specific binding sites was performed using 5% (w/v) nonfat dry milk in TBST for 1hr at room temperature followed by three 5-minute washes in TBST. Membranes were incubated with primary antibody solutions (using 5% milk or BSA in TBST) overnight at 4°C at the respective dilutions: E-Cadherin (BD, #610181,1:5000), Vimentin (Abcam, ab92547, 1:500), CD44 (CST, #3570, 1:200), Zeb (CST, #3396, 1:200), Slug (CST, #9585, 1:200), Twist (GeneTex, #GTX127310, 1:500), GAPDH (CST, #5174, 1:10000). Membranes were then washed three times in TBST, 5 minutes each time and incubated with appropriate secondary antibodies in blocking solution for 1hr at room temperature. Following three 5-minute washes with TBST, immunoreactivity was visualized using a standard ECL detection system.

### Phase-Contrast and Confocal Fluorescence Microscopy

Phase-contrast images of cells undergoing EMT were obtained with a Zeiss Axiovert 40C microscope (5x, 10x, 20x objectives) equipped with an Axiocam 105 color camera and processed using the Zen 2 software. For confocal fluorescence microscopy, HCC827 cells that were previously grown in either control conditions (absence of TGFβ, 2% FBS RPMI media) or in presence of TGFβ (5ng/ml) for a number of days were seeded on coverslips placed in 6-well plates at a density of 125 000 cells per well. Treatment with TGFβ was continued depending on time-course requirements. Following treatment, cells were rinsed with HBSS and subsequently with ice cold 5% BSA/PBS twice. Cells were then fixed with paraformaldehyde solution (PFA, EMS, #15710) at a final concentration of 4% for 15 minutes at room temperature followed by 3 washes with PBS. Permeabilization was performed with 0.1% Triton x100 (Sigma Aldrich) for 10 minutes at room temperature followed by 3 washes with PBS. Blocking was performed by incubating cells with 5% BSA/PBS solution for 30 minutes at room temperature. Cells were then incubated with fluorophore conjugated primary antibodies for 1 hr in the dark (Alexa 488 Vimentin Antibody (BD, #562338) and Alexa 647 E-Cadherin Antibody (BD, # 324112)). Following 3 washes with PBS, cells were mounted in Gold antifade reagent with DAPI (Molecular Probes P36935). Images were obtained and analyzed using the inverted Zeiss LSM 880 laser scanning confocal microscope with Airyscan (40x objective) and Zen Black software respectively.

### Flow Cytometry

Following treatments cells were lifted off tissue culture plates using TrypLE (Life technologies, #12605-010). After counting and assessing % viability with Trypan Blue exclusion, 1 x 10^6^ cell aliquots per condition were fixed by adding PFA at a final concentration of 1.6% for 10 minutes at room temperature. Cells were then centrifuged at 500g for 5 minutes at 4°C to pellet cells and remove PFA and washed once with cell staining media (CSM, 0.5% w/v BSA, 0.02% w/v NaN_3_ in PBS). Cells were permeabilized with methanol solution for 10 minutes on ice and optionally stored at −80°C for long-term storage. After two washes with CSM, master mix of antibodies was added to pelleted cells at a total volume of 100uL for 30 minutes in the dark at room temperature. Following two washes with CSM, cells were analyzed using a LSR II.UV. Cell viability was assessed using the LIVE/DEAD Fixable Blue Dead Cell Stain Kit (Life Technologies, #L23105) as per manufacturer’s instructions. Fluorophore conjugated primary antibodies used: PE/Cy7 E-Cadherin (Biolegend, Clone 67A4, #324115), Alexa 488 Vimentin (BD, Clone RV202, #562338), Pacific Blue CD44 (Biolegend, Clone IM7, #103019), APC/Cy7 CD24 (Biolegend, Clone ML5, #311131), Alexa 647 Twist (Bioss, #bs-2441R). LSR II. UV shown were created in Cytobank (www.cytobank.org, Cytobank, Inc., Menlo Park, CA) as previously described ^54^.

### Tumor Dissociation and Processing

Briefly, fresh specimens were immersed and transferred from Stanford Hospital to the laboratory in MACS Tissue Storage Solution (Miltenyi, #130-095-929). After recording tumor weight, obtaining macroscopic pictures and removing fat and necrotic areas, tumors were cut into pieces of 2-4 mm. Tumor dissociation was performed utilizing the MACS Tumor Tissue Dissociation kit (Miltenyi, #130-095-929) per manufacturer’s instructions. Tumor derived single-cell suspensions were centrifuged at 500g for 5 minutes to pellet tumor cells which were subsequently resuspended in RPMI and applied on a MACS SmartStrainer (70uM, Miltenyi #130093237,) for filtration. Following centrifugation (500g, 5 minutes) red blood cells were removed using the Red Blood Cell Lysis Solution according to manufacturer’s instructions (Miltenyi, #130094183). After tumor cells were washed and re suspended with RPMI, we exposed them to cisplatin (Sigma-Aldrich #P4394, final concentration 5µM) for 5 minutes at room temperature to label dead cells for subsequent mass cytometry analysis. Complete RPMI media was added to quench cisplatin reaction and tumor cells were washed twice. Cell viability was determined with trypan blue staining prior fixation (to ensure high viability before processing). All samples analyzed were > 80% viable. Finally, cell aliquots of 0.5-1 x 10^6^ cells /mL were fixed with PFA at a final concentration of 1.6% for 10 minutes at room temperature and washed twice with CSM before being stored in CSM at −80°C until barcoding and staining procedures for CyTOF analysis.

### Mass Cytometry Antibodies

Antibodies used for mass cytometry analysis, including respective information on antibody clone, vendor, metal isotope and staining concentration, are summarized in Table S1. Aside from one antibody purchased from Fluidigm (CD104), all antibodies were conjugated to metal-isotopes in-house using the MaxPar Antibody Conjugation Kit (Fluidigm) and titrated to determine optimal staining concentrations.

### Sample Processing, Mass-Tag Barcoding and Antibody Staining for Mass Cytometry Analysis

Cell line samples were processed similar to tumor samples following dissociation and as previously described^54^. Briefly, following treatments, cells were lifted off tissue culture plates using TrypLE. Cell count and initial % viability were assessed using Trypan Blue exclusion. To assess cell viability for mass cytometry, cell pellets were briefly incubated in 1mL PBS containing cisplatin (final concentration 5 µM) for 5 minutes at room temperature. Cisplatin reaction was quenched by adding 2ml of CSM and subsequent centrifugation for 5 minutes at 500g. Cell pellets were resuspended in CSM in 0.5-1×10^6^ aliquots per condition and were fixed by adding PFA at a final concentration of 1.6% for 10 minutes at room temperature. Cells were centrifuged at 500g for 5 minutes at 4°C to pellet cells and remove PFA and washed once with CSM. Cell pellets were resuspended in CSM and stored at −80°C until all time-points of the same time-course or multiple clinical specimens were collected. To improve staining consistency, samples were palladium barcoded and pooled for staining as previously described^55^. In short, different combinatorial mixtures of palladium-containing mass-tag barcoding reagents in DMSO were added to each cell line or tumor sample previously resuspended in PBS-saponin solution and mixed with pipetting. Samples were incubated with barcoding reagents for 15 minutes at room temperature. Reaction was quenched with the addition of CSM followed by several washes with CSM prior to pooling all samples together to proceed with staining. Surface and intracellular antigen antibodies were separately added into two master mixes in CSM, filtered through a 0.1 µm filter (Millipore #UFC30VV00) and centrifuged at 1000g for 5 minutes to remove antibody aggregates. For tumor specimen analysis, separate staining cocktails using the same concentrations were prepared with the addition of antibodies towards CD45, FAP and CD31 for gating out immune, stromal and endothelial tumor populations respectively as previously described ^13^. Samples were first incubated with surface antibody master mix for 30 minutes at room temperature. Cell samples were then washed with CSM and permeabilized with methanol for 10 minutes on ice. Following 2 washes with CSM, samples were then incubated with the intracellular antibody master mix for 30 minutes at room temperature. Samples were washed twice with CSM and resuspended in PBS containing 1:5000 191Ir/193Ir MaxPar Nucleic Acid Intercalator (Fluidigm) and 1.6% PFA to stain DNA and stored at 4°C for 1-3 days. Prior to mass cytometry analysis, cells were washed once with CSM and twice with filtered double-distilled water that was previously resuspended in normalization beads (EQ Beads, Fluidigm), filtered and kept on ice. During event acquisition, pooled samples were kept on ice at all times and introduced into the CyTOF 2 (Fluidigm) using the Super Sampler (Victorian Airship and Scientific Apparatus, Alamo, CA, USA). Apart from antibody metal isotopes listed in Table S1, we also recorded event length, barcoding channels (102Pd, 104Pd, 105Pd, 106Pd, 108Pd, 110Pd), normalization beads (140Ce, 151Eu, 153Eu, 165Ho, 175Lu), DNA (191Ir and 193Ir), and dead cells (195Pt and 196Pt).

### Mass Cytometry Data Processing

Normalization and single-cell debarcoding were performed through respective algorithms as described previously^55^. Debarcoded samples were uploaded as separate FCS files and analyzed on Cytobank. Cytobank software was used for traditional cytometry statistics and visualization (histograms, density plots, heatmaps and SPADE (as previously described^56^).

## QUANTIFICATION AND STATISTICAL ANALYSIS

### CCAST Clustering

Raw mass cytometry data from cell line TGFβ time-course experiments were arsinh transformed on Cytobank. After gating out dead and apoptotic cells, remaining cells from all time-points were pooled together. The CCAST-recursive partitioning-based algorithm^24^ was applied assuming a minimum of 8 clusters on a subsample of ∼96000 cells obtained from density-dependent down sampling ^56^ based on 6 EMT markers (E-Cadherin, Vimentin, CD44, CD24, MUC1 and Twist). These markers were selected using an unbiased non-parametric regression tree analysis^57^. All 6 markers were among the most statistically significant (p-value <0.001) markers that correlated independently with the top 3 principal components explaining about 50% variability in the data (See Table S2). Final selection of the 6 markers was based on EMT biological relevance. The CCAST analysis resulted initially in 13 clusters (see Supplementary Fig. 4) of which 5 were excluded from downstream analysis based on a pre-set threshold of each cluster ≥1% of total number of cells analyzed. All data processing and clustering analysis was performed in R.

### Force-Directed Layout Visualization of CCAST Clusters

To visualize the spatiotemporal dynamics of EMT and MET processes in HCC827 cells treated with TGFβ, we used Vortex, a graphical tool for visualizing clusters generated from multiparametric datasets. Specifically, we created single-cell force-directed layouts. The resulting graph was constructed specifically on the pre-identified 8 CCAST clusters. We utilized Vortex to only implement edge connections between subsequent time points (similar to the FLOW-MAP algorithm^58^). The force-directed layout graph was used to assess the EMT/MET phenotypic continuum in terms of timing and the expression profiles of all 27 markers measured with mass cytometry. The graph is built by repulsing all cells with forces proportional to how dissimilar they are in multidimensional protein expression space, while edges hold adjacent cells together by constant spring-like force ^26^.

### Construction of the 2D EMT-MET STAte MaP (STAMP)

Using the 6 CCAST clustering markers (E-cadherin, Vimentin, Twist, CD24, CD44, and MUC1), we next generated a 2D-t-SNE^59^ map using the Rtsne package implemented in R with a default perplexity parameter of 30. We next applied Voronoi mapping using 8 optimal center points obtained from the above CCAST clusters. These points were obtained from cluster-specific bins with sizes estimated by varying sizes of 1 to 8 units over time and selecting the size showing the largest adjacent change in the number of cells. To place a continuous boundary on the t-SNE map we next applied a generalized Convex Hull mapping algorithm, which combines a Voronoi diagram and Delaunay triangulation^60^ using the alphahull R package. The combined t-SNE-Voronoi mapping-Convex Hull analysis produces a 2D EMT/MET state space partition map capturing the plasticity of EMT and MET processes.

### Neural Network for Projecting onto the EMT-MET STAMP

To model the non-linear structure of the underlying 2D EMT-MET state map, we constructed a single-layer-hidden artificial neural network model using the above 6 markers as input and the reduced 2D t-SNE feature space as a bivariate response. Specifically, we used the 6 input expression values for each cell from the cell line samples to predict the corresponding position of each new cell onto the 2D reference map space. Projection results were visualized as bivariate scatter plots. To train the network, weights on the edges are modified to minimize the error in the output. At the end of training, the inputs which are most important in prediction have the largest weights while those that are less important have lower weights. The network was trained on 90% min-max normalized data using the “nnet” R package and the remaining 10% was used (1) to optimize for the 11 hidden nodes needed and (2) to carry out a 10-fold cross validation on the stability of model predictions. The trained neural network was next used to project an independent HCC827 time-course repeat, A549 and H3255 cell line samples and 5 clinical specimens. A k-nearest neighbor classification on the partition centers was carried out for each sample to estimate the densities of cells in various EMT state partitions on the map.

### Inferring EMT and MET Trajectories from Time-Course Data: TRAjectorty of CElls Reconstruction (TRACER) algorithm

To more rigorously test the hypothesis of hysteresis when comparing EMT and MET, we developed a novel trajectory reconstruction analysis TRACER, that does not rely on pseudotime assumptions and allows branching. We assumed that EMT and MET are classical Markov processes^33^ with constant transition probabilities between the states within EMT and MET but can differ between EMT and MET. Because our observations of the proportion of cells in each state are made at discrete, nonuniform time points, and because we do not observe the trajectories of individual cells, we cannot employ standard methods to estimate for the state transition probabilities^34^. Also, because the number of states is relatively large compared to the number of observations, we employed sparsity assumptions^35^. We modeled the state transition probabilities *p*_*jkt*_ between states *j* and *k* at time *t* by imposing sparsity penalties on *p*_*jk*t_ for *j* not equal to *k*, but no penalty on *p*_*jjt*_, with the idea is that there is no cost for staying in a state, but switching between states is discouraged. Under these assumptions, the transition probabilities EMT and MET can be estimated by convex optimization, with the sparsity parameter λ selected through cross-validation. We chose by the 1-standard error rule^61^, that is, we chose the largest λ such that its error is within 1-standard error of the error of the minimizing. This leads to more parsimonious solution. Using bootstrap analysis, we provide a distribution of the transition probabilities for EMT and MET, as shown in Fig.4E. For the bootstrap analysis, we first independently sampled a multinomial distribution for cell counts in each state. We split the bootstrap sample into 2 folds. The first fold was used to estimate the sparsity parameter λ. We evaluated the estimator with the chosen λ on the second fold. For representative EMT and MET networks, the medoid networks for EMT and MET was selected among all the bootstrap samples as the network for which the average distance to all other networks (entry-wise L1 distance between transition matrices) was smallest (Fig. 4F).

## DATA AND SOFTWARE AVAILABILITY

Mass cytometry time-course data will be made available through Cytobank.

*Flow Cytometry analysis and confocal imaging for this project was done on instruments in the Stanford Shared FACS and Cell Sciences Imaging Facilities.*

