## Supplementary figures and tables for "Mapping Lung Cancer Epithelial-Mesenchymal Transition States and Trajectories with Single-Cell Resolution"

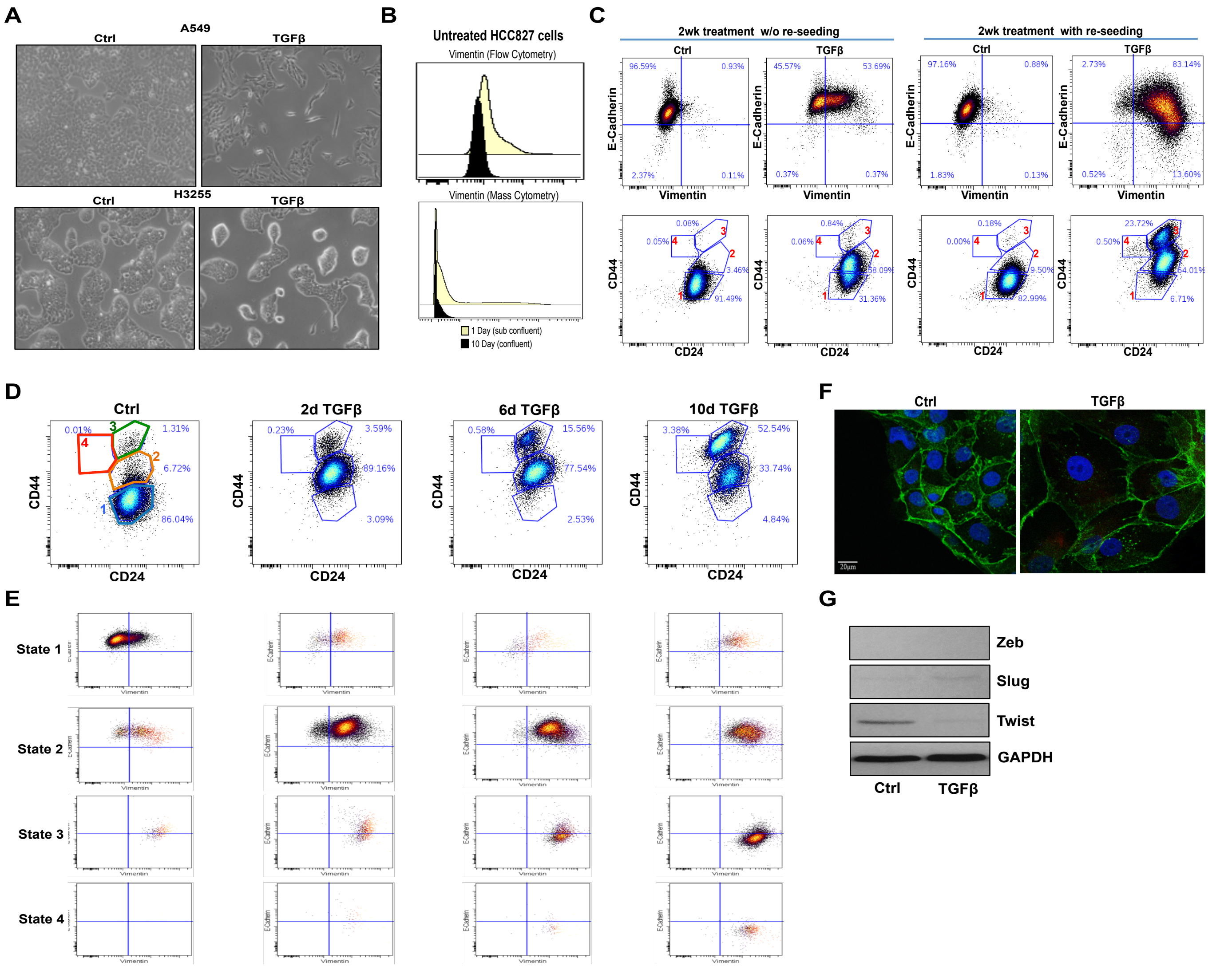

**Supplementary Figure 1. Optimization of TGFβ-induced EMT in NSCLC cell lines and identification of canonical EMT states via flow cytometry**

(A) Representative images of A549 and H3255 cells treated with TGFβ (5ng/mL) for 1 and 2 weeks respectively.
(B) Vimentin levels in untreated HCC827 cells that were in either sub confluent (yellow color, seeded and collected after 1 day) or confluent (black color, seeded and collected after 10 days) cell culture conditions. Top graph, flow cytometry analysis, bottom graph, mass cytometry analysis of an independent biological replicate experiment.
(C) E-Cadherin/Vimentin and CD44/CD24 flow cytometry plots of HCC827 cells treated with TGFβ for 2wks in non-re-seeding (left) and re-seeding cell culture conditions (right). Shown are the respective 4 canonical EMT states and % of cells per gated area.
(D) CD44/CD24 flow cytometry plots shown in Figure 1A, with the respective 4 gated states and % of cells per gated area
(E) Gated states 1 through 4, shown independently towards their E-Cadherin/Vimentin phenotype expression.
(F) Representative confocal images of HCC827 cells treated with TGFβ (5ng/mL) for 6 days and stained for E-Cadherin (green). Note the significantly larger cell size of cells treated with TGFβ compared to ctrl cells, while still retaining E-Cadherin expression at tight junctions.
(G) Immunoblots of EMT transcription factors Zeb, Slug and Twist in HCC827 cells treated with TGFβ for 2 weeks.

| Antibody Target | Clone | Vendor | Metal | Mass | Final Concentration (ug/mL) |
|---|---|---|---|---|---|
| CD45 | H130 | Biolegend | Yb | 89 | 1 |
| FAP | F11-24 | eBioscience | Ln | 113 | 1 |
| CD44 | IM7 | Biolegend | Ln | 115 | 1 |
| cleaved Caspase 3* | C92-605 | BD | Nd | 142 | 1 |
| phospho-Src | K98-37 | BD | Nd | 144 | 1 |
| phospho-EGFR | D745 | CST | Nd | 145 | 2 |
| EGFR | D38B1 | CST | Nd | 146 | 1 |
| TROP2 | 77220 | R&D | Nd | 148 | 3 |
| Oct 3/4 | O50-808 | BD | Nd | 150 | 4 |
| Notch3 | MHN3-21 | Biolegend | Eu | 151 | 1 |
| Cytokeratin 8 | SP102 | Abcam | Sm | 152 | 1 |
| PD-L1 | 29E.2A3 | Biolegend | Eu | 153 | 2 |
| MUC1 | SPM492 | Abcam | Sm | 154 | 2 |
| RUNX1 | 1C5B16 | Biolegend | Gd | 155 | 2 |
| Snail | C15D3 | CST | Gd | 156 | 4 |
| E-Cadherin | 67A4 | Biolegend | Gd | 158 | 2 |
| Nanog | polyclonal | CST | Tb | 159 | 2 |
| phospho-H3 | HTA28 | Biolegend | Gd | 160 | 2 |
| CD24 | ML5 | Biolegend | Dy | 161 | 4 |
| phospho-SMAD2/3 | D27F4 | CST | Dy | 162 | 2 |
| phospho-NFKb | K10-895.12.50 | BD | Dy | 163 | 2 |
| phospho-S6 | N7-548 | BD | Dy | 164 | 2 |
| phopsho-Rb | J112-906 | BD | Ho | 165 | 0.5 |
| Cytokeratin 7 | SP52 | Abcam | Er | 167 | 0.5 |
| Twist | polyclonal | Bioss | Er | 168 | 4 |
| non phospho-b-catenin | D13A1 | CST | Er | 170 | 1 |
| CD31 | WM59 | Biolegend | Yb | 171 | 0.5 |
| Slug | C19G7 | CST | Yb | 172 | 4 |
| CD104 | 58XB4 | Fluidigm | Yb | 173 | 2 |
| Vimentin | D21H3 | CST | Yb | 174 | 2 |
| phospho-AMPK | 40H9 | CST | Lu | 175 | 2 |

* Replaced in certain runs with cleaved PARP antibody (Clone: F21-852, Vendor: BD, Pr 141, 1ug/mL)

**Supplementary Table 1. Mass Cytometry (CyTOF) EMT-MET antibody panel. Related to Figure 2**
Antibodies in grey boxes were included in CyTOF runs of NSCLC clinical specimens to separate immune (CD45), stromal (FAP) and endothelial (CD31) cell populations from tumor epithelial cells (see Methods for additional information).

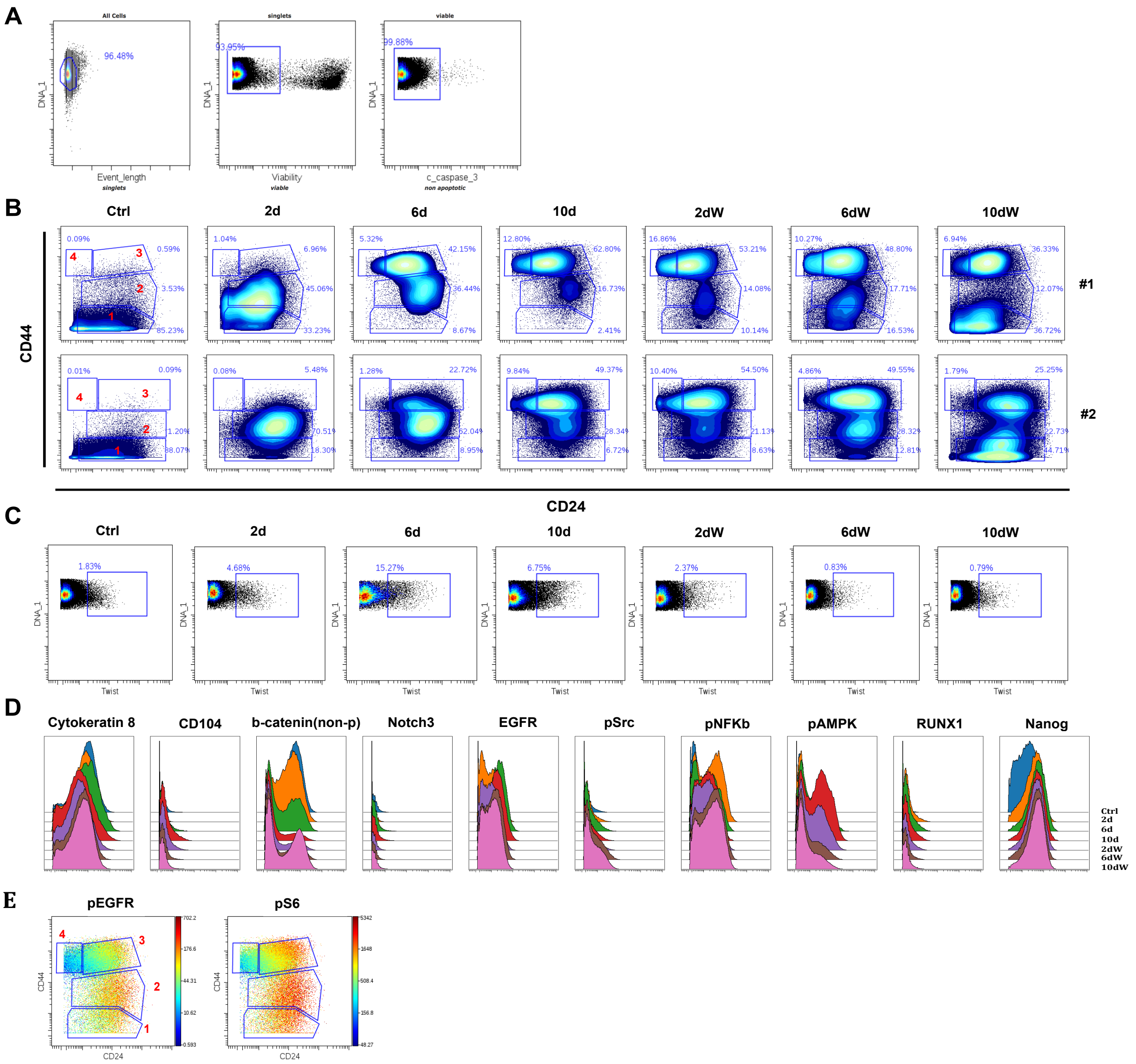


**Supplementary Figure 2. Time-course analysis of EMT and MET by Mass Cytometry. Related to Figure 2**

(A) Gating strategy shown for a HCC827 time-point sample analyzed with mass cytometry. Following de-barcoding, hand-gating of DNA and cell length parameters were used to separate single cells from debris. Dead and apoptotic cells were excluded using positive cisplatin and cleaved caspase-3 staining respectively.

(B) Mass cytometry measurements of CD44/CD24 expression changes confirm the existence of the 4 canonical EMT states observed in HCC827 cells with flow cytometry analysis in two independent biological TGFβ time-course replicates. Shown also are % of cells in each of the 4 states and how these change with time.

(C) Gating Twist positive cells in HCC827 time-point samples (depicted in Figure 3) shows the transitional increase in numbers during EMT (4 and 6d TGFβ) and subsequent decrease prior time-point in which the majority of cells become most mesenchymal (10d TGFβ and 2dW).

(D) Remaining cellular markers measured with mass cytometry in the HCC827 TGFβ time-course depicted in Figure 2.

(E) State 4 ($CD44^{hi}$/$CD24^{lo}$) cells are negative for pEGFR and pS6 expression.

| PCA1 | PCA2 | PCA3 |
|---|---|---|
| pRb | **CD44** | **Twist** |
| pEGFR | **Vimentin** | Cytokeratin7 |
| Nanog | **ECadherin** | pEGFR |
| Cytokeratin8 | pAMPK | pS6 |
| pS6 | Oct3/4 | |
| pNFKB | TROP2 | |
| **CD24** | Slug | |
| pSmad2/3 | **MUC1** | |
| **ECadherin** | | |

**Supplementary Table 2. Statistically significant markers resulting from principal component analysis (PCA) of mass cytometry data. Related to Figure 3.**
In bold are the 6 EMT markers selected among all the markers for our CCAST analysis. E-Cadherin, Vimentin, CD44, CD24, MUC1 and Twist were among the most statistically significant (p-value <0.001) markers that correlated independently with the top 3 principal components which explained about 50% variability in the data (see Methods for additional information).

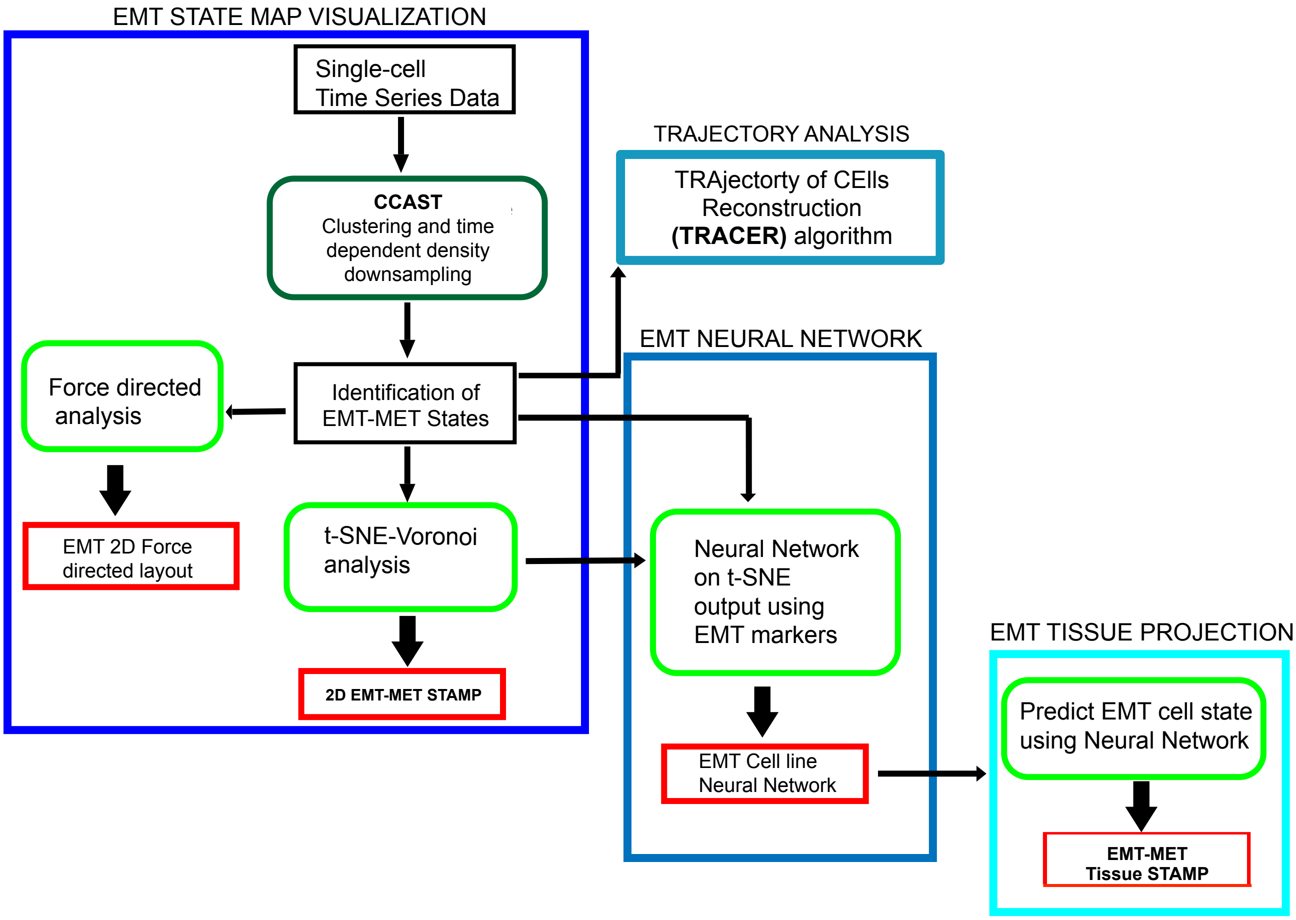


**Supplementary Figure 3. Flowchart depicting computational analyses and tools applied and/or developed for mapping EMT-MET states and trajectories. Related to Figures 3-6. See Methods for details**

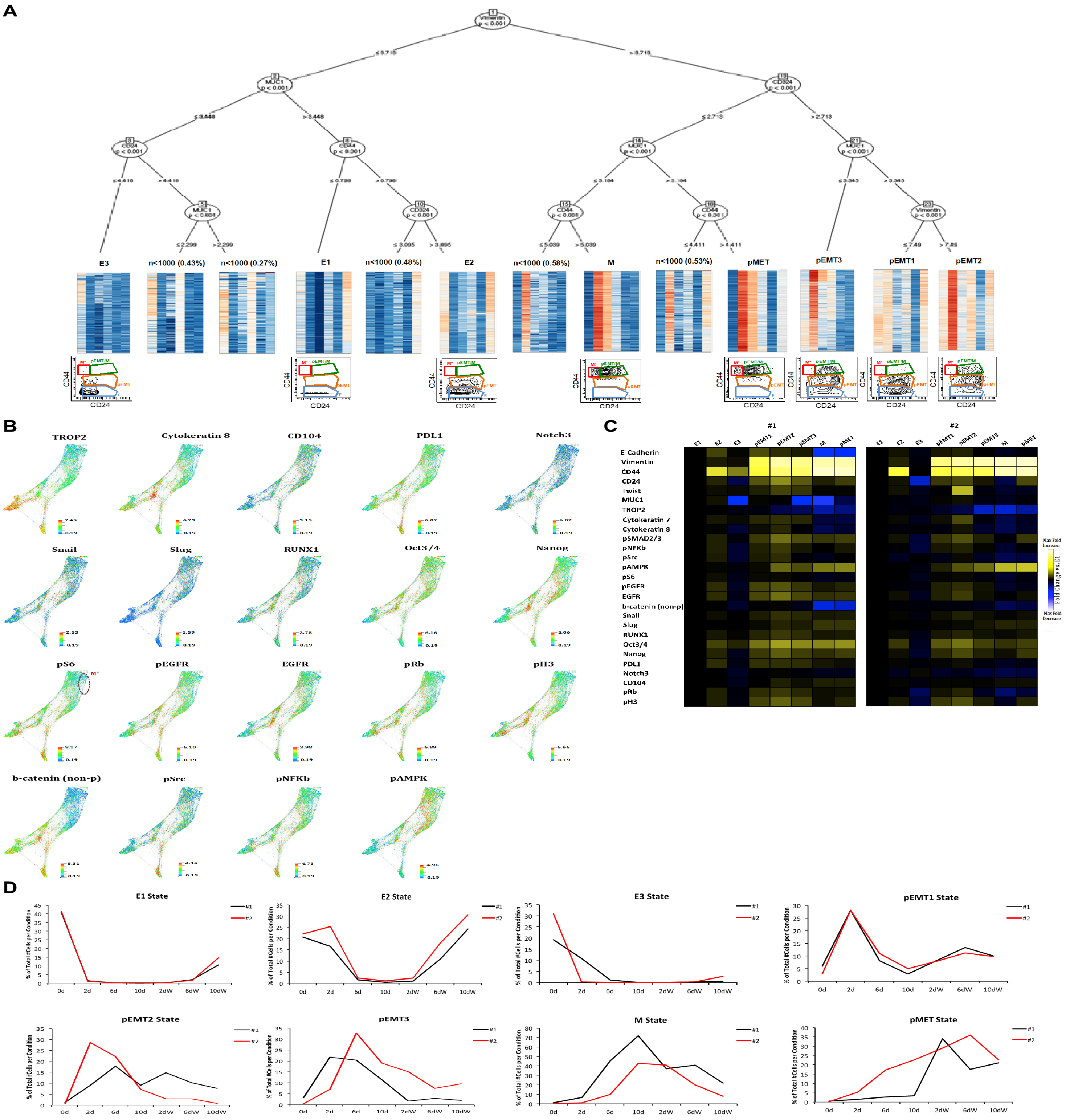


**Supplementary Figure 4. Time-course analysis of EMT and MET by Mass Cytometry. Related to Figure 3**

(A) CCAST decision tree for the HCC827 mass cytometry data derived from all 28 markers showing 13 distinct subpopulations using 6 clustering markers: E-Cadherin, Vimentin, CD44, CD24, MUC1 and Twist. Eight of the 13 clusters met our criteria (n>1000 cells, ≥1% of pooled cells) for downstream analysis. (*Bottom*) Heat maps of normalized data from all subgroups derived from the decision tree above and their expression for the aforementioned 6 clustering markers. Below each computationally derived EMT-MET state is the respective CD44/CD24 plot showing how it relates to the EMT canonical states.

(B) Force-directed layouts (FDLs) colored by protein expression levels of the remaining markers analyzed with mass cytometry. Note red circled area on the pS6 FDL, indicating M$^*$, CD44$^{hi}$/CD24$^{lo}$ subpopulation of cells.

(C) Heat map summary depicting fold change expression of each marker per EMT/MET state towards E1 state from experiment shown in Figure 3 (#1) and in an independent biological replicate experiment (#2).

(D) EMT/MET state dynamics shown for experiment in Figure 3B (black lines, #1) alongside matched states in an independent biological replicate experiment (red lines, #2)

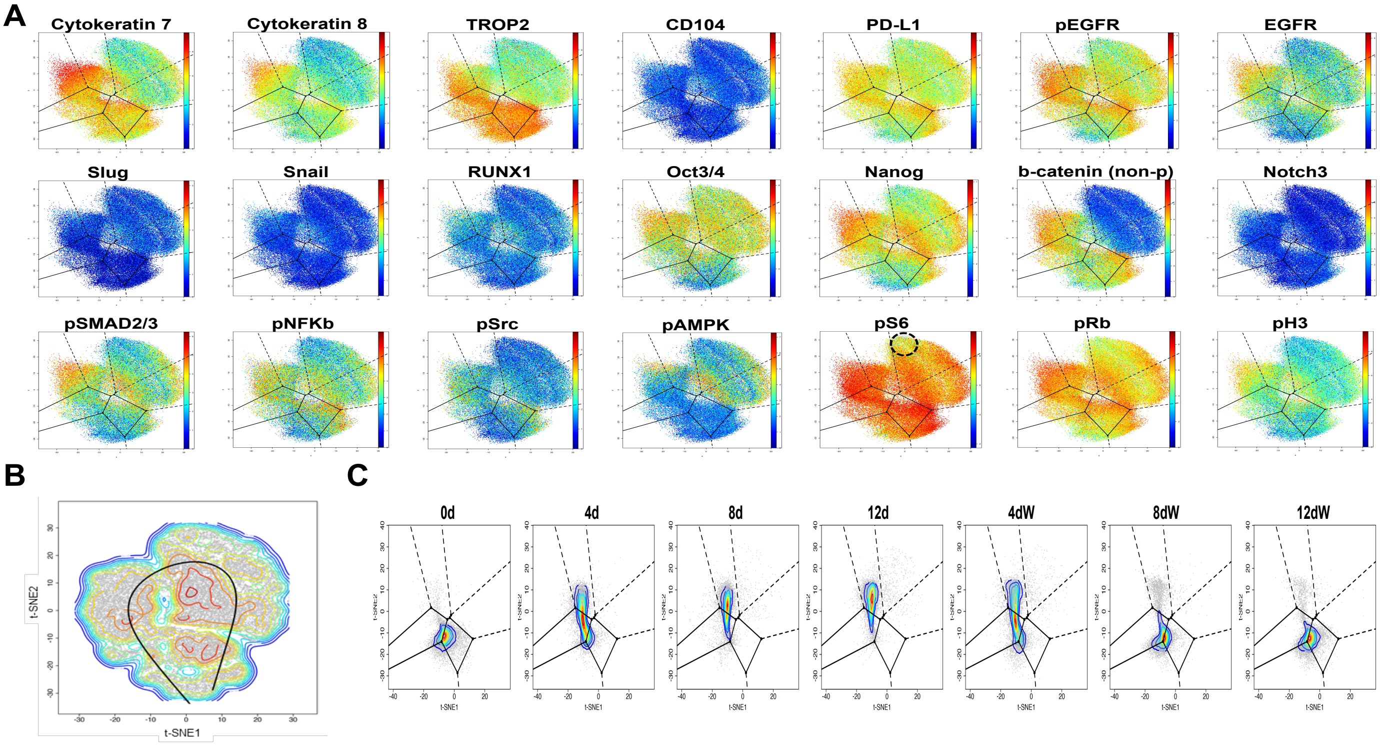


**Supplementary Figure 5. Construction of an EMT-MET STAte MaP (STAMP). Related to Figure 4**

(A) Expression profiles of remaining markers in pooled HCC827 time-point data visualized on the EMT/MET state map. Circled areas on pS6 plot depicts the M*, CD44$^{hi}$/CD24$^{lo}$ subpopulation of cells.

(B) Slingshot analysis confirms the trajectory that involves the *pMET* state. *pMET* is visited by cells during TGFβ withdrawal conditions and is unique to one of two possible MET scenarios.

(C) Time-point t-SNE density plots of HCC827 cells that did not efficiently undergo EMT during a TGFβ time-course experiment. Note that at all time-points a very small number of cells occupy the M region and almost no cells occupy the pMET region of the map.

| Case No. | Description | Grade (Differentiation) | Size (cm) | Mutations | Smoking History |
|---|---|---|---|---|---|
| 1 | NSCLC, Adenocarcinoma | Well | 4.7 | EGFR (exon 19 deletion) | Never |
| 1 | NSCLC, Adenocarcinoma | Moderate | 4 | EGFR (p.Leu858Arg) | Former |
| 2 | NSCLC, Adenocarcinoma | Poor | 2.5 | EGFR (exon 19 deletion) TP53 (p.Asp259Val) | Never |
| 3 | NSCLC, Adenocarcinoma | Poor | 5 | TP53 (p.Arg248Leu) | Former |
| 4 | NSCLC, Adenocarcinoma | Moderate/Poor | 3.5 | KRAS (p.Gly12Ala) | Former |

**Supplementary Table 3. Clinical data of the 5 patient specimens analyzed with mass cytometry**

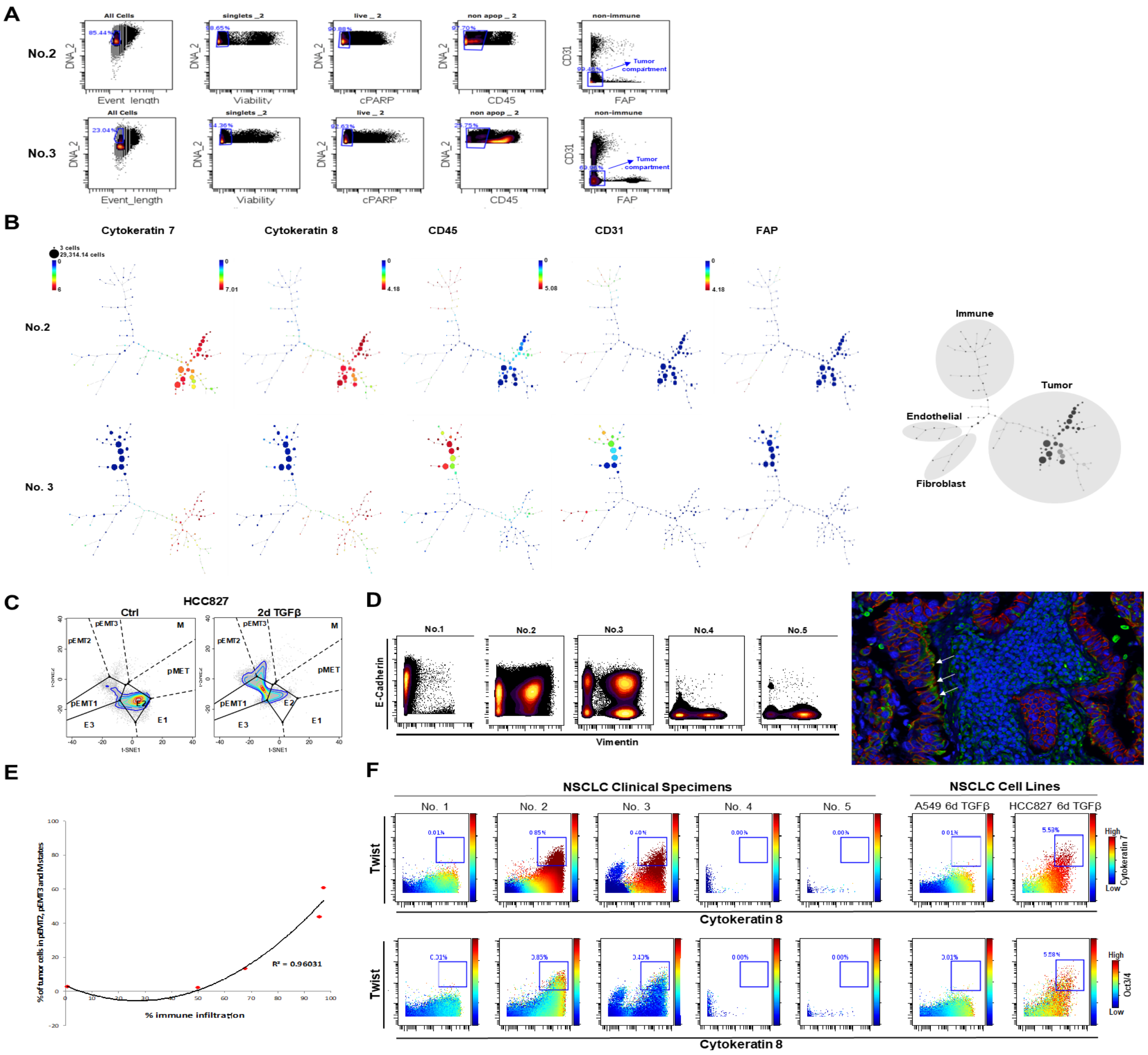

**Supplementary Figure 6. Mass cytometry analysis and projection of NSCLC clinical specimens onto the EMT-MET STAMP. Related to Figure 6**

(A) Gating strategy for the computational separation of tumor cells for two NSCLC clinical specimens (No.2 and 3). Viable, non-apoptotic single cells were separated as described in Methods. Further gating out of immune, endothelial and stromal cells was performed with CD45, CD31 and FAP antibodies respectively.

(B) SPADE analysis showing efficient computational separation of tumor cells with the use of CD45, CD31 and FAP antibodies in clinical specimens No.2 and 3. Cytokeratin 7 and 8 expression profiles are shown to confirm absence in immune, endothelial and stromal populations, and variable expression in tumor cells. Staining profiles of all markers (CD45, CD31, FAP, Cytokeratins 7 and 8) can be used to identify and gate out cell populations in clinical specimens analyzed with mass cytometry (SPADE illustration of clinical specimen No.2 to the right, shaded areas).

(C) Projection of HCC827 cells (ctrl, 2-day TGFβ) that were stained and analyzed alongside clinical specimens for validation purposes.

(D) E-Cadherin/Vimentin mass cytometry plots of the 5 NSCLC clinical specimens that were analyzed (left). Immunofluorescent staining of matched tissue from specimen No.2 (right). Arrows indicate pEMT cells co-expressing E-Cadherin (red) and Vimentin (green).

(E) % of tumor cells projected on pEMT2, pEMT3 and M areas of the EMT-MAP map vs. % immune infiltration shows positive correlation between the two in the five NSCLC clinical specimens analyzed.

(F) Twist$^+$, Cytokeratin 7$^+$, Cytokeratin 8$^+$, Oct3/4$^+$ subpopulation of cells is detected in EGFR mutated NSCLC (Clinical Specimens No. 2 and 3)

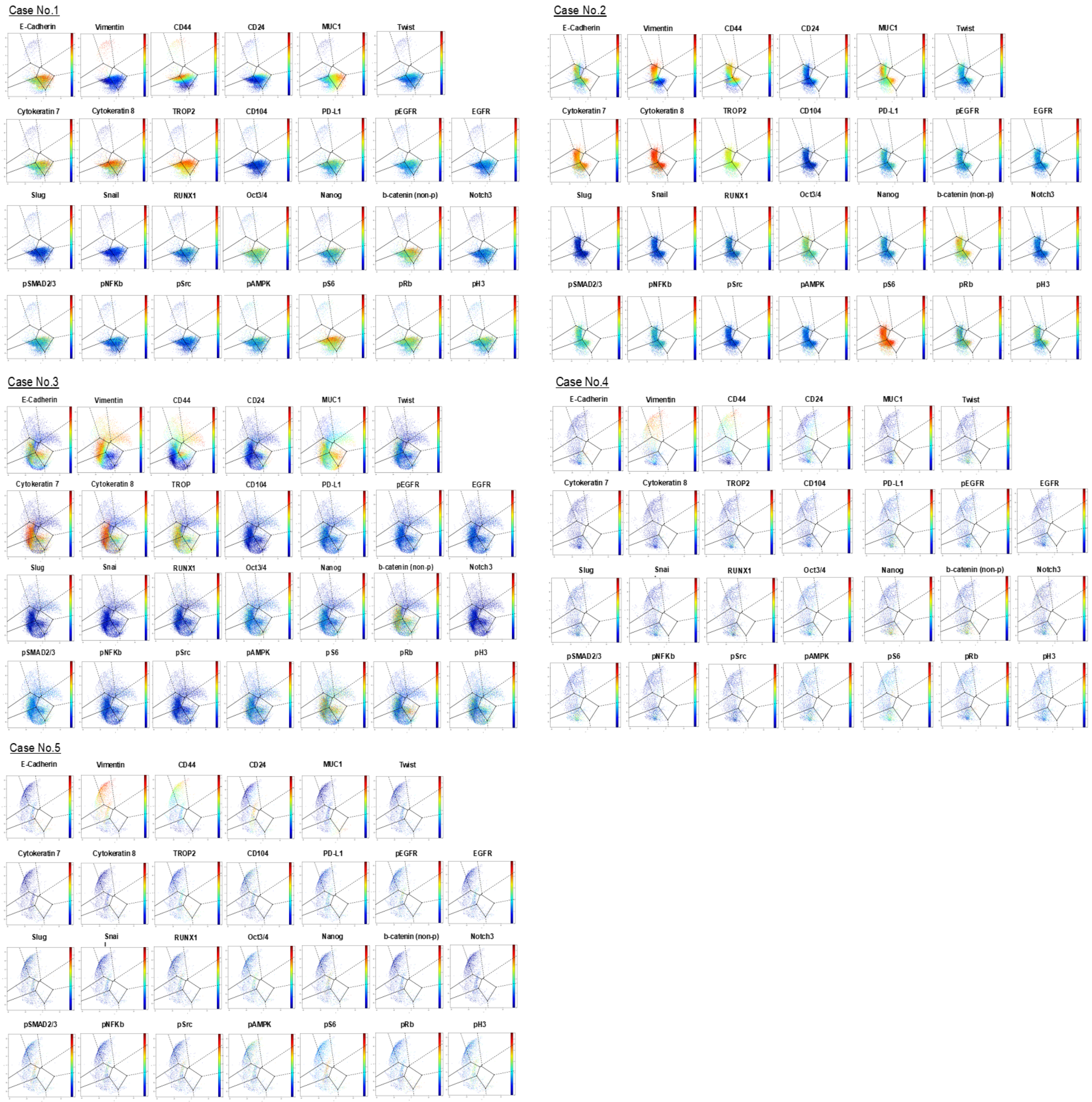


**Supplementary Figure 7. Expression profiles of all markers in the NSCLC clinical specimens visualized on the EMT-MET STAMP. Related to Figure 6**

Clinical specimens No.1, 2, and 3 all share EGFR mutations and map primarily on E and pEMT1 regions of the map, with the exception of specimen No.3 which carries an additional TP53 mutation, and has cells mapping on more mesenchymal regions of the map including those of pEMT2, pEMT3, M and pMET. Clinical specimens No. 4 and 5 carry mutations other than EGFR, specifically TP53 and KRAS respectively, and have a proportion of cells mapping on the mesenchymal pEMT2 and M regions of the map. Of note, specimens No. 4 and 5 had the highest % immune infiltration at 95.85% and 97.20% respectively.
